# DNA co-methylation networks outline the structure and remodeling dynamics of colorectal cancer epigenome

**DOI:** 10.1101/428730

**Authors:** Izaskun Mallona, Susanna Aussó, Anna Díez-Villanueva, Víctor Moreno, Miguel A. Peinado

## Abstract

Epigenomic plasticity is interconnected with chromatin structure and gene regulation. In tumor progression, orchestrated remodeling of genome organization accompanies the acquisition of malignant properties. DNA methylation, a key epigenetic mark extensively altered in cancer, is also linked to genome architecture and function. Based on this association, we postulate that the dissection of long-range co-methylation structure unveils cancer cell’s genome architecture remodeling.

We applied network-modeling of DNA methylation co-variation in two colon cancer cohorts and found abundant and consistent transchromosomal structures in both normal and tumor tissue. Normal-tumor comparison indicated substantial remodeling of the epigenome covariation and revealed novel genomic compartments with a unique signature of DNA methylation rank inversion.

## Introduction

Cancer cell functional reprogramming involves gene expression dysregulation driven by genetic and epigenetic changes. The contribution of epigenetic mechanisms to malignant phenotypes has been thoroughly studied and includes extensive DNA methylation alterations as prominent features of most cancers types (Portela and Esteller, 2010; Feinberg et al., 2016). DNA methylation mainly occurs in the cytosine of the CpG dinucleotide and is usually associated with a repressed chromatin state. Changes in DNA methylation have multiple effects in genome regulation and have been directly associated with gene overexpression and silencing, chromatin remodeling and chromosomal instability (Eden et al., 2003; Rodriguez et al., 2006; Jones, 2012; Schübeler, 2015; Feinberg et al., 2016). Direct comparison of the DNA methylation profiles in the tumor versus the paired normal tissue reveals both losses (hypomethylation) and gains (hypermethylations) of the epigenetic mark. The extent of the change may range from discrete sites and promoters to large regions (Frigola et al., 2006; Portela and Esteller, 2010; Hansen et al., 2011; Jones, 2012; Feinberg et al., 2016).

Neighboring CpGs have a higher chance of being similarly methylated (Eckhardt et al., 2006; Shoemaker et al., 2010; Libertini et al., 2016); nonetheless, the actual extent of this vicinity effect is disputed, with reports of complete to weak or very low decay of co-methylation as the genomic distance increases in different cell types and tissues (Li et al., 2010; Akulenko and Helms, 2013; Fortin and Hansen, 2015; Salhab et al., 2018). Most studies about the functional impact of DNA methylation changes have focused the analysis to local effects on neighboring genes (Jones, 2012; Schübeler, 2015). More recently, taking advantage of the availability of genome-scale DNA methylation data from large cancer datasets, the study of DNA co-methylation profiles has been addressed from different points of view, including the analysis of long range correlations (Akulenko and Helms, 2013; Fortin and Hansen, 2015; Zhang and Huang, 2017) and gene centered approaches (Li et al., 2014; Gao and Teschendorff, 2016; Saghafinia et al., 2018). Numerous methods have been developed to evaluate differential methylation and differential variability of DNA methylation (Jaffe et al., 2011; Jenkinson et al., 2017; Teschendorff and Relton, 2018; Jenkinson et al., 2018; Rulands et al., 2018; Visakh and Nazeer, 2019; Libertini et al., 2018; Jenkinson et al., 2019).

We hypothesize that dynamically methylated elements reveal the functional organization of human cancer cell’s genome. To get insights into the structure, functional determinants and underlying mechanisms of DNA methylation dynamics we examined the DNA methylomes of colon cancer patients by evaluating loci with shared patterns of DNA methylation. The rationale was that samples subjected to complex physiopathological processes (e.g. tumor initiation and progression), albeit being highly heterogeneous, share common driver and passenger events from which biologically relevant phenotypic traits arise.

## Methods

We separately run our analysis using two independent colon cancer datasets. Our primary cohort, Colonomics (http://www.colonomics.org), included 90 paired primary tumors (stage IIA and IIB) and their adjacent normal tissue. Of the 90 patients, 67 were males and 23 females, aged 43-86 years (mean: 70.37), and 20 developed metastasis. All tumors were microsatellite-stable. Samples were evaluated for DNA methylation (Illumina Infinium HumanMethylation450 BeadChip Array) and gene expression (Affymetrix Human Genome U219), and somatic mutations (exome sequencing) (Closa et al., 2014; Cordero et al., 2014; Sanz-Pamplona et al., 2015; Sole et al., 2014).

The Cancer Genome Atlas (TCGA) series was composed by 256 primary tumor and 38 adjacent nontumor samples from the colon adenocarcinoma (COAD) cohort (Zhu et al., 2014). Patients were aged 31 to 90 years at diagnosis (mean 65.61), and included 141 males, 144 females and one unassigned. Pathologic stages included Stage I (40), Stage II (97), Stage III (75), Stage IV (32); 11 were not available or discrepant. Regarding microsatellite instability, 10 were positive, 65 negative and 181 were either not tested or had an unknown status. Samples readouts included DNA methylation by Illumina Infinium HumanMethylation450 BeadChip Array, gene expression by RNA-Seq counts and somatic mutations by exome sequencing.

Data processing and workflow is depicted in Figure 1. Briefly, DNA methylation beta values were subjected to serial pairwise correlation analysis for the Colonomics tumor series (primary dataset) and normal adjacent tissue, as well and both TCGA normal samples and tumors (external datasets). Strong associations (effect size Spearman’s *ρ* ≥ 0.8) were stored. Co-methylation networks were built upon the correlations data (Cullen and Frey, 1999; Csardi and Nepusz, 2006; Clauset et al., 2009; Gillespie, 2014; Delignette-Muller and Dutang, 2015), from which highly connected modules according to the fast greedy community detection algorithm were isolated (Csardi and Nepusz, 2006; Clauset et al., 2004) were isolated. Purity correction was estimated as in (Zheng et al., 2017; Aran et al., 2015). Next, modules were functionally annotated by comparing them to the literature (Lister et al., 2009; Hansen et al., 2011; Aryee et al., 2014; Fortin and Hansen, 2015) and molecular features databases (Liberzon et al., 2011; Heinz et al., 2010; Smyth, 2005; Quinlan and Hall, 2010; Love et al., 2014; Gel et al., 2015). Modules characterization, including reproducibility assessment, consisted in mutual profiles comparison and differential expression analysis among different cohorts (Hubert and Arabie, 1985; Shannon et al., 2003; Krzywinski et al., 2009; Langfelder et al., 2011; Akdemir and Chin, 2015).

**Figure 1:**
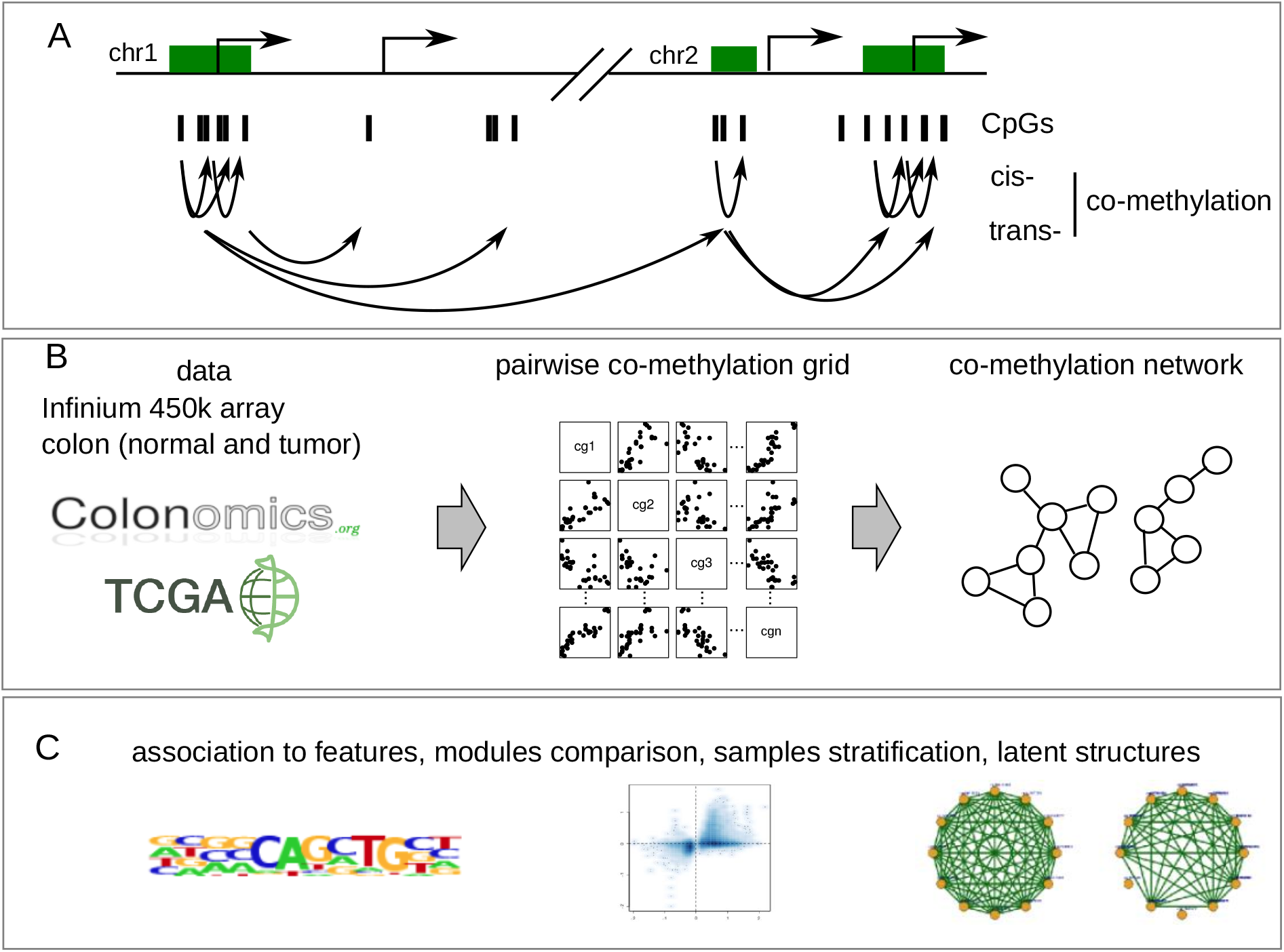
Co-methylation analysis framework. A, CpGs co-methylation occurs at close (cis) and long distances (trans). B, DNA co-methylation networks describe the distance-agnostic co-methylation structure in colon tumors and normals. C, co-methylation networks associate to molecular signatures.

We implemented a Web service at http://maplab.cat/corre to facilitate browsing the DNA co-methylation events of any locus or gene. The tool allows candidate queries either by gene symbol or Illumina Infinium probename, providing the annotated co-methylations full list. Apart of downloadable spreadsheets, Corre renders interactive plots to evaluate zonal (chromosome) and functional (chromatin color) enrichments (Ernst et al., 2011; Gesmann and de Castillo, 2011; Zhang et al., 2013; Conway et al., 2016). Source code is available at https://bitbucket.org/imallona/corre under the GPL terms. Corre can be accessed freely and without registration.

Extended methods are available as supplementary material.

## Results and discussion

### Close and distant CpGs co-methylate in colon cancer samples

We retrieved DNA methylation data as measured by Infinium HumanMethylation450 Array *β* values from 90 tumor and 90 adjacent normal tissues (Closa et al., 2014; Cordero et al., 2014; Sanz-Pamplona et al., 2015; Sole et al., 2014) to feed the co-methylation analysis (Figure 1). Quality check consisted in three steps: First, we excluded probes non uniquely mapping to a genomic location, being polymorphic or located in sex chromosomes (Price et al., 2013). Second, probes with low variability (standard deviation *s*_*β*_ < 0.05) were filtered out to get rid of correlations led by outliers, while enriching for CpGs with intermediate methylation (Du et al., 2010). Finally, probes with missing data in any sample were removed. Next we sequentially calculated the bulk pairwise Spearman’s correlations between any possible pair of probes adjusting for multiple testing (Table S1).

Co-methylations were frequent at neighboring CpGs but also detectable at long distances (Figure 2A), being this effect independent of local probe density (Figure S1). The extent of the comethylation as measured from the correlation coefficient *ρ* (ranging from −1 to 1, in which 0 means full independence) were bell-shape distributed, thus indicating that the majority of correlations lied on the non-significant range, as expected, and with a tendence towards positive association at CpGs located at short distances due to local co-methylation. Opposite methylation trends are detectable in trans (Figure 2). Coordinated variation was independent of the CpG location in open or closed chromatin compartments (Figure S2).

**Figure 2:**
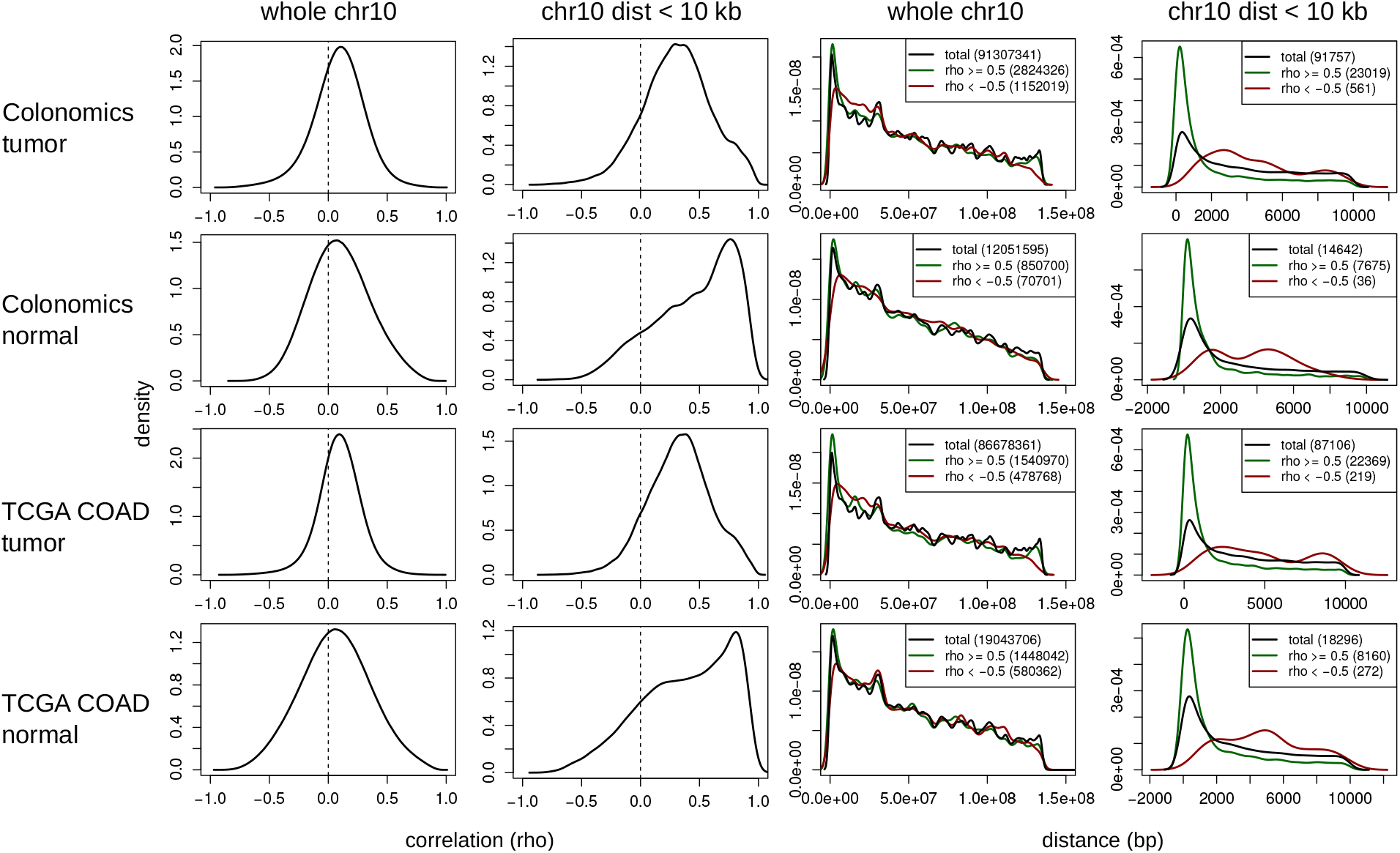
Co-methylation and distance effects. Correlations distribution is bell-shaped and shifted towards positive values at the tumor cohort (whole chr10); the trend to co-methylate is noticeably increased in cis (chr10 probes located at less than 10 kbp). Red: negative correlations (*ρ* < − 0.5); green: positive correlations (*ρ* ≥ 0.5).

To underpin the biological relevance of the findings and rule out the co-methylation structure arising due to technical noise, we evaluated five possible sources of artifacts: multiple testing, batch effects, leading outliers, tumor purity and chip design and none of them appeared to have a significant effect on the results.

To account for the iterative nature of the analysis, consisting in exhaustively computing any pairwise correlation between the Infinium probes with variable DNA methylation, we set an astringent effect size cut-off of the Spearman’s correlation cofficient *ρ* ≥ 0.8, which is close to the conservative Bonferroni p-adjustment for the datasets used (optimization against the asymptotic p-values as calculated by the Fisher Z transform, Table S1) (Fisher, 1915; Shakhbazov et al., 2016). The absence of notorious clustering of DNA methylation values (Figure S3) indicates that batch effects are unlikely drivers of co-methylation (Leek et al., 2010). To attenuate the leading effect of DNA methylation outliers we filtered in probes with sufficient variation in DNA methylation (setting a standard deviation threshold). The number of variable probes was about twice in the tumor series as compared to normal tissue (Table S1). To reinforce robustness against outliers, nonparametric Spearman correlations were run, which rely on DNA methylation ranks rather than values (Croux and Dehon, 2010). Regarding tumor purity, we found co-methylation modules to retain tight correlation structure after subtracting purity effects (Zheng et al., 2017) (Figure S4). As for the Infinium array design, neither the probes GC content (Figure S5) nor the dye channel (Figure S6) drove the correlations structure; probes mapping to multiple locations or overlapping to SNPs were filtered out (Price et al., 2013).

### Anatomy of the distance-agnostic co-methylation network in colorectal cancers

Given the pairwise nature of co-methylation events, we envisioned a co-methylation network linking all significant co-methylations in a genomic location-agnostic manner, thus representing genome-wide comethylation trends, and potentially linking shared drivers. To do this, we selected the top-scoring correlations and assembled a network in which loci and correlations are represented by nodes (vertex) and links (edges). For the sake of simplicity, we only considered edges with positive correlations.

The tumor network resulted in 63,130 nodes and 26 million co-methylations (Table S2). The distribution of each CpG degree (amount of connectivity, number of co-methylating neighbors) showed a heavy-tail shape with the vast majority of nodes being linked to few counterparts, whereas a few nodes displayed a fairly abundant connectivity. The degree distribution did not resemble power-law, lognormal nor exponential, as there was not linear dependency between the cumulative frequencies and the connectivities (Figures S7 and S8). This structure was unaffected by loci features such as chromatin state (Figure S9) and genomic category (Figure S10); but lost when filtering out trans interactions, as probes placed at any distance in cis showed power-law compatible distributions (Figure S11), as described previously (Zhang et al., 2017).

Interestingly, genomic loci acting as co-methylation hubs (frequently co-methylating in trans) presented homogeneous intermediate DNA methylation levels in normal samples (Figures S12 and S13), which have been suggested to play regulatory roles (Elliott et al., 2015), with an important enrichment of imprinted loci (n=53, 11%). On the other side, they were not enriched for partially methylated domains (PMDs), which have been reported as loci with intermediate DNA methylation values and high variability (Lister et al., 2009): 27% (132 CpGs) of the rich probes overlapped PMDs, similarly to the 33% (21,065 CpGs) of the probes with at least a significant co-methylation and what is expected from the background of the whole set of Infinium probes, with an overlap of the 31% (147,257 CpGs).

### The colon cancer co-methylation landscape is reproducible across cohorts and tumor type-specific

To test the reproducibility of the network we repeated the analysis using an independent dataset, the COAD cohort from TCGA, consisting of 256 primary colon adenocarcinomas. In TCGA, the DNA methylation *β* value calling procedure differs from the original dataset, and therefore reduces the chance of covariation artifacts arising due to the data processing bias.

Near a quarter million probes fulfilled the variability criteria in TCGA colon tumors (Table S1). The overall correlations distribution and the co-methylation decay with distance matched that of the primary dataset (Figures 4 and 2). Next, we evaluated whether the correlation value for each pair of probes was conserved, including the non-significant pairs. To do so, we computed exhaustive pairwise correlations of CpGs located at the chromosome 10 against itself and plotted the original *ρ* values of each CpG pair against validation dataset from TCGA. The linearity of both landscapes (Figure 4) indicated a high concordance of the overall co-methylation levels. Similarly, the TCGA colon cancer co-methylome network was comparable in terms of graph properites (Table S2 and Figure S7). Interestingly, the comethylation landscape was very dissimilar to other tumor types’ (Figure S14), whilst it was comparing colon datasets from the starting and the validation cohorts, strongly supporting the specificity of the colon cancer co-methylation network.

In a similar vein and to compare the structure of both networks, we checked whether the correlating nodes present in both cohorts displayed the same connectivity to other nodes. The influence score of each node (e.g. based in the number of links arising from it) was estimated using the PageRank score (Page et al., 1999) in both datasets. Both primary and validation tumor co-methylation networks showed a reproducible distribution of nodes’ PageRanks mostly composed by lowly influential CpGs (Figure S15).

### DNA co-variation in normals pinpoints massive co-methylation rearrangements in tumors

We wondered whether some of the co-methylations found in cancers were already detectable in adjacent normal colonic mucosa, and to which extent the co-methylome structure differed from the tumor. The normal colon co-methylome network was built using 90 non-tumor tissues (Table S1). We found that both the number of probes fulfilling the variance prerequisite (n=99,346) and the number of total correlations (7,430,741) decreased to 39% and 22% of the tumor’s ones, respectively. This result was consistent with the higher DNA methylation variability in tumors. As expected, a predominance of positive correlations was observed, being more intense for close probes (Figure 2A). Anti-methylations showed much more abundant in tumors than in normal tissue (Figure 2). In agreement with the associations found in tumors, co-methylations were underrepresented in active promoters (Figure S16) and the co-methylation network’ connectivities were not power-law distributed (Figure S7). Probes pairs correlation values showed partial agreement between the normal and tumor datasets (Figure 4). Interestingly, age and other demographic features, common to both the tumor and adjacent normal data, were not primary drivers of the co-methylation (Figure S17).

In order to assess the reproducibility of the network, we repeated the analysis using normal samples from the external dataset from TCGA. It should be noted that this dataset only includes 38 normal samples, and as the correlation significance depends on the sample size (Fisher, 1915), keeping the same cut-offs is likely to boost the number of false positives. On the other hand, increasing the *ρ* cut-off to an equivalent detection threshold (*ρ* = 0.96, Table S1) produced a very small network whose properties might be out of scale with the previous analysis. With this cautionary note in mind, keeping the *ρ* = 0.8 cut-off the network confirmed the distinctive distribution of pairwise correlations (Figure 4), whose differences are especially conspicuous at short ranges (Figure 2).

### Co-variation in DNA methylation segments the epigenome into sparse compartments

In order to dig into the observed trend of preferential co-methylation of some loci, we explored whether the network had highly connected subnetworks (also known as modules or communities). Modules consist of clusters of nodes heavily interconnected as compared to the rest of the network ers of nodes heavily inter-connected (as compared to the rest of the network) constitute a community (Newman and Girvan, 2004; Fortunato, 2010). Modularity is quantified as the fraction of edges connecting nodes of the same type minus what it is expected in a randomly wired network. Scores of 0 indicate no modularity and networks with modular structure typically range from 0.3 to 0.7 (Newman and Girvan, 2004). The tumor co-methylation network was found to be modular (modularity = 0.47) (Table S2), and we partitioned it into 3,270 modules ranging from two to 18,727 nodes using the Clauset, Newman and Moore’s fast greedy method (Clauset et al., 2004). Interestingly, the normal tissue network exhibited a higher co-methylome modularity (0.62) (Table S2) and network segmentation resulted in 1,265 modules ranging from two to 17,758 nodes. To rule out the effect of tumor purity on correlation structure, we regressed it out and found that co-methylation modules retained tight correlation structure (Zheng et al., 2017) (Figure S4).

Expectedly, we found the vast majority of the small modules to be composed by sets of probes located at close distance from each other (e.g. at CpG islands). As these could be mined by standard block variation or differential methylation analysis, we focused in trans-chromosomal modules, which were more numerous in the tumor co-methylome (1% of the total modules) than in the normal’s (1.4%). Trans modules are diverse in physical properties, such as the chromosome, and in functional properties, such as A/B compartment or chromatin state (Figure S20). In agreement with the original dataset results, TCGA tumor co-methylation network was also modular (modularity score 0.41) (Table S2) and segmentation produced 3,421 modules ranging from two to 8,981 nodes. The application of size and distance filters reduced the number of trans-chromosomal modules to 35.

The validation series TCGA similarly depicted a modular tumor network (modularity score 0.41) (Table S2) and its clustering into modules resulted in a segmentation in 3,421 modules ranging from two to 8,981 nodes. By application of the same size and co-location filters applied to the original series (at least 10 probes, 1 Mb apart or in different chromosomes) only 35 modules remained. Next, we faced both cohorts module by module in order to detect assess their preservation.

We next evaluated the extent of conservation of transchromosomal co-methylation territories between normals and tumors (biological effect) as well as between cohorts (replicability). Networks clustering on adjusted Rand’s distance (Hubert and Arabie, 1985) indicated that modules memberships separate tumor’s from normal’s networks in both datasets (Figure S21A), in line with the similarities in nodes population (Figure S21B), and their spatial co-methylation patterns (Figure S22). It is worth noting that the use of a correlation threshold (i.e. effect size *ρ* ≥ 0.8) may underestimate module co-methylation maintenance when the correlations distribution gets displaced towards values below, but close to, the statistical significance cut-off (Figure S23).

Module preservation across tissue types and cohorts was also evaluated by cross-tabulation of the number of shared CpGs. Twelve tumor modules had one or more counterparts in the normal tissue network, and a similar number in the validation TCGA tumor cohort (Fisher’s exact test) (Figures S24 and S25). The five-top sized Colonomics tumor modules partially matched to multiple TCGA’s modules. This result might be technically due to the resolution limit of modularity-optimizing algorithms to detect modules, which tend to aggregate modules into few giant components, disregarding their inner complexity (Fortunato and Barthelemy, 2007).

### A novel and sparse genomic compartment shows inverted DNA methylation ranks in colon tumors

As we detected a noticeable gain of anti-methylations in the tumors as compared to the normals (Figure 2), and that the co-methylation tumor network depicted two separate giant components (Figure 3) that were not present in the normal tissue, we suspected a major restructuring of co-methylation architecture. The external series from TCGA replicated this finding in both tumor and normal epigenomes (Figure 3).

**Figure 3:**
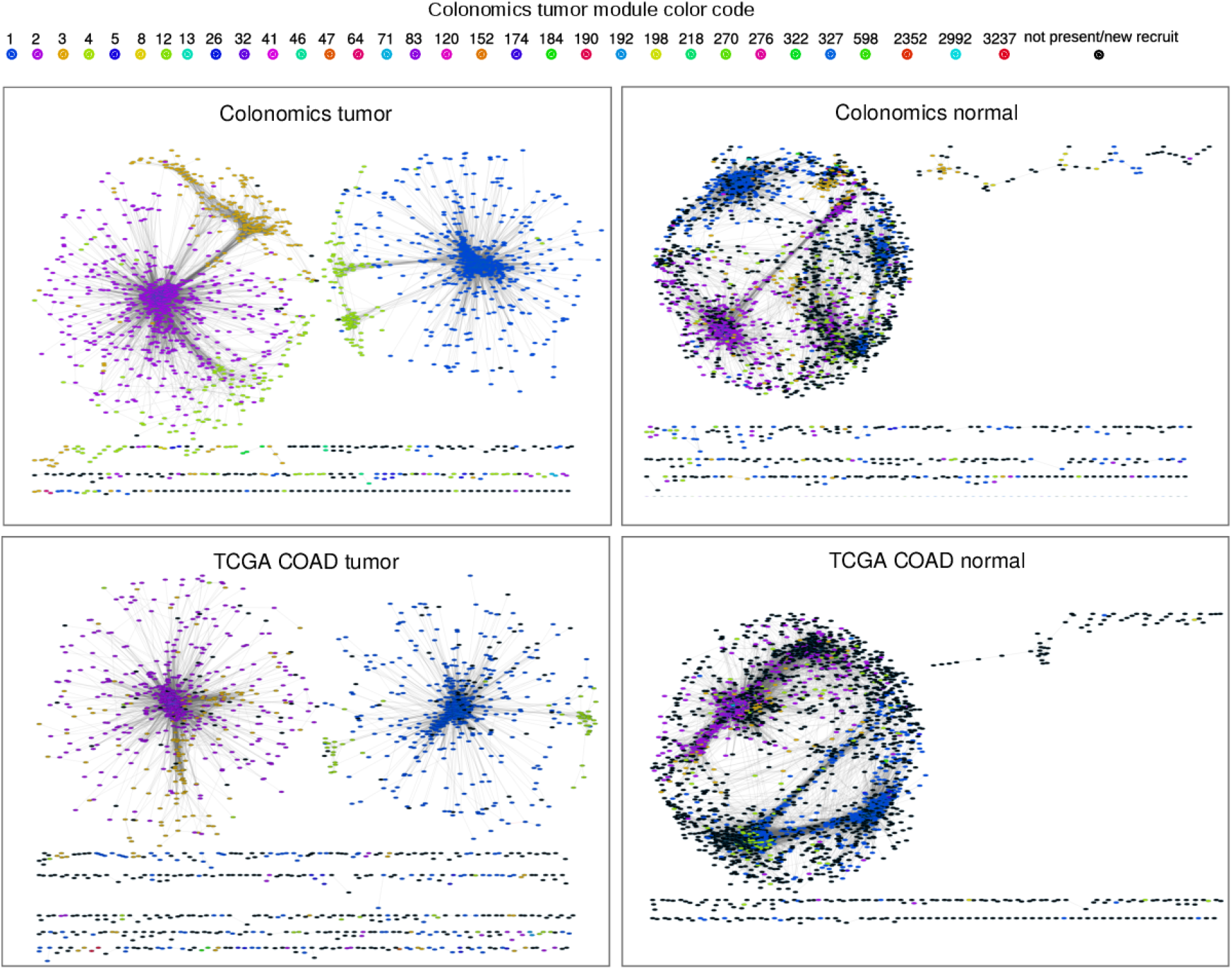
Co-methylation network module memberships comparison. Networks, built independently for each dataset, highlight the nodes (CpGs) by the trans module they belong to at Colonomics tumor (black indicates new recruits or, for Colonomics tumor, non-trans module membership). Graphs are limited to a random sample of 5000 nodes and solitary nodes are not plotted; network layout was calculated by 1 − *ρ* (edges) weighted springs.

**Figure 4:**
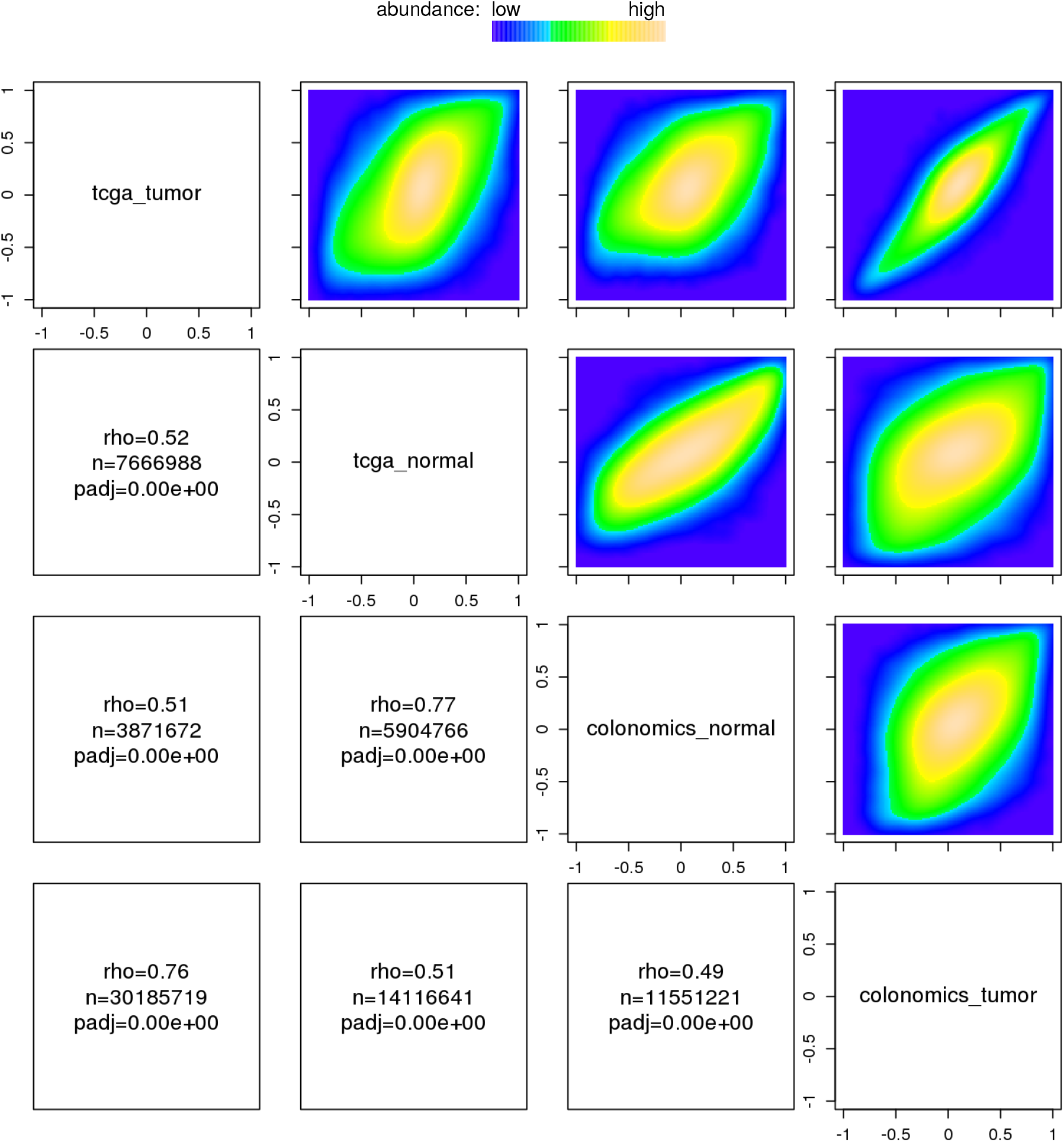
Correlations replication across cohorts. For each cohort, we conducted an exhaustive pairwise correlation analysis for any probe with at least *sd* ≥ 0.05. Then we pairwise plotted the correlation coefficient rho (X and Y axis) for each probe pair to check whether the co-methylation landscape was reproduced. Analysis was restricted to chr10.

The pervasive nature of anticorrelations overcame age, gender, tumor stage, tumor purity and anatomical site potential effects on modules’ DNA methylation variation (Figures S17, S18, S19). Loci with copy number alterations also conveyed the module-specific DNA methylation ranks mimicking the profiles along balanced regions (Figure S26).

This observation was caused by the inversion of the co-variation pattern of sparse co-methylation compartments, whose DNA methylation status gets a quantitative DNA methylation rank inversion during malignant transformation (Figure 5). To describe this phenomenon, which displays properties beyond regional differential methylation nor increased regional DNA methylation variability, we next evaluated the anatomical and functional features of the co-methylation modules.

**Figure 5:**
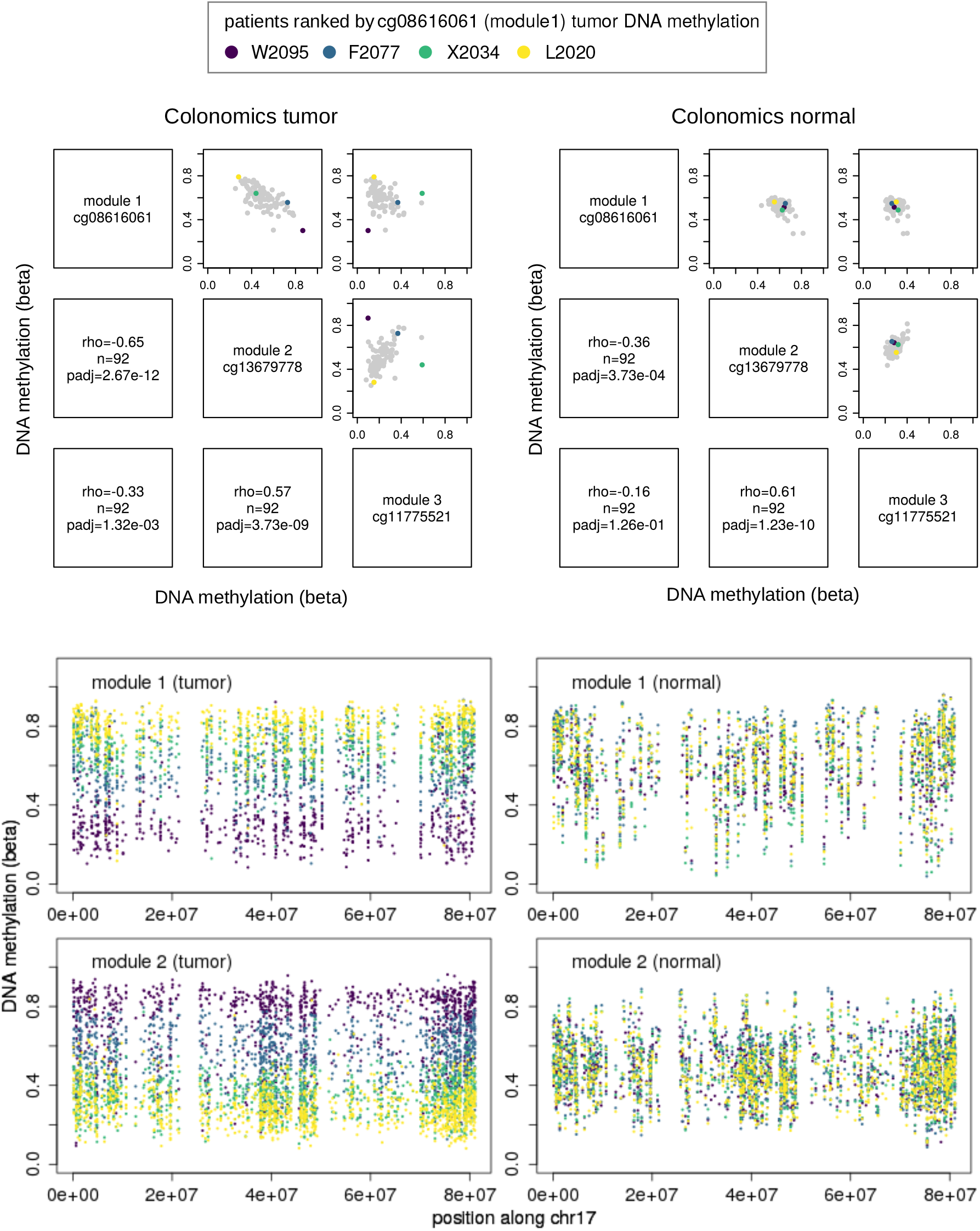
Opposed DNA methylation ranks. Tumor samples show opposed DNA methylation ranks at module 1 and module 2. Colored dots depict four patients sorted according to their tumor DNA methylation values at module 1.

### Co-methylation loci are enriched for functional signatures

To test the hypothesis that co-methylation structures are directly related with functional properties we investigated genomic and functional features of co-methylated CpGs. We subtracted the overrepresentation of promoters and promoter-related features in the Infinium array (Bibikova et al., 2011; Sandoval et al., 2011; Silva-Martínez et al., 2017), as well as their presumable effects driven by spatial clustering (Kim et al., 2008; Barrera and Peinado, 2012; Gaidatzis et al., 2014; MacDonald et al., 2015; Libertini et al., 2016; Wang et al., 2016). The tumor modules displayed important differences in feature enrichment, including chromatin states, genomic categories, CpG islands and association with known motifs (Figure 3). In line with the weighted gene co-expression network analysis, in which functional signatures can be told apart by mining gene co-expression (Langfelder and Horvath, 2008), the module-specific co-regulation patterns denoted by distinctive abundance of genomic and functional states. Among the multiple features analyzed, a striking global enrichment of inactive promoters was observed in a large number of modules (Figure S27), pointing out potential clusters of co-regulated genes.

Next, we explored the co-methylation modules overlap with regions of DNA methylation variability previously reported in colon cancer, including differentially methylated regions (Hansen et al., 2011). Interestingly, seven out of 32 tumor modules significantly overlapped tumor hypermethylated blocks (Figure S28). Regarding other types of DNA methylation variability reported by Hansen et al. (2011), modules showed distinctive profiles, with frequent enrichment in boundary shifts as well in loss of regulation; novel hypomethylation blocks were enriched in three modules only. This complexity reinforces the individuality of co-methylation modules.

A comprehensive summary of structural and functional feature enrichment for each co-methylation module is shown in Tables S4 and S5). To name a few examples, multiple co-methylation modules were significantly enriched for Polycomb-related marks (i.e. H3K27me3, or SUZ12, EED and PRC2 targets; e.g. tumor modules 1, 3, 5 and 598); for frequently mutated at COSMIC molecular signatures (i.e. tumor modules 2, 4 and 8); and for gene expression (i.e. tumor module 8). Finally, we could also confirm that co-methylation network associated features found in TCGA matched Colonomics enrichment signatures, e.g., the underrepresentation of co-methylations within active promoters (Figure S27).

To shed light into causal factors driving dynamic methylome modularity we searched for enriched motifs (i.e. transcription factor binding sites) at the co-methylating loci (Supplementary methods). We found that six out of the 32 Colonomics tumor modules presented one or more significantly enriched motifs (Table S6). Enriched motifs included ETS and RUNX families in modules 1, 3, 327, FOS family members (FRA1, ATF3, BATF, FOSL2, JUN-AP1) in modules 2 and 4), FOXA1-related factors (FOXA1, HNF4a, FOXMA) in module 2, GC box (KLF5 and KLF4) in module 2, C/EBP in module 3, PAX7 and MYF5 in module 5, homeobox in modules 5 and 83, MADS in module 152, and ASCL1 in module 598.

In summary, the modularity of long range DNA co-methylation events in colorectal cancer point to novel properties of epigenetic remodeling. We postulate that coordinated DNA methylation changes at interspersed sites (here identified as belonging to the same module) might be linked to common players and effectors, hypothesis in line with the DNA methylation-mediated binding of transcription factors to regulatory sites (Kribelbauer et al., 2017; Yin et al., 2017).

### A colon cancer co-methylome browser

We developed a web tool available at http://maplab.cat/corre to ease browsing of raw and processed data. To illustrate the usage of this tool we queried the INHBB gene encoding activin B, a member of the TGF-beta family, with different biological activities, including a role in cell proliferation and inflammation. Epigenetic silencing of INHBB is frequent in colorectal cancer (Frigola et al., 2006) and has been proposed as indicator of poor outcome (Mayor et al., 2009). The co-methylation landscape of INHBB exposed by the Corre tool showed a large number of positive correlations 20 kb upstream and downstream of the gene in both normal and tumor samples (Figure S29). Negative correlations were only present in tumors and were enriched in poised promoters, indicating the potential remodeling of bivalent states and hypermethylation (McGarvey et al., 2008; Ohm et al., 2007; Rodriguez et al., 2008). Compared with the normal samples, the tumors displayed an increase in the number of links for most sites, although some chromosomes, especially 8 (Figure S29E), but also 13, 14, 17 and 21, showed an opposite trend with a depletion of co-methylations in the tumors as compared with the normal samples (Figure S30). Another interesting result was in regard to cg03699182 probe (Figure S29D, arrowhead) located in the CpG island of the INHBB promoter presented 42 co-methylations in the normal samples against only three in the tumors. Most of the cg03699182 co-methylations affected were located in poised promoters of polycomb regulated genes (Figure S30). The dynamics of the connections and the properties of the affected sites are consistent with the participation of instructive mechanisms resulting in the DNA hypermethylation and long range epigenetic silencing of multiple genes in colorectal cancer (Frigola et al., 2006; Keshet et al., 2006; Michieletto et al., 2017).

## Conclusions

Here we describe a network modeling analysis of DNA methylation co-variation. When applied to colorectal cancer and normal colonic mucosa it is demonstrated that the architecture of DNA methylation variability is modular and embodies structural and functional associations with a potential impact in cancer biology. Normal and tumor networks display characteristic and differential structures that are consistently replicated in independent cohorts. Interestingly a quantitative inversion of DNA methylation ranks results in a novel and sparse genomic compartment pinpointing a long-range functional reorganization of cancer cell’s epigenome.

## Ethics approval and consent to participate

The study protocol was approved by the Clinical Research Ethics Committee (CEIC) of the Bellvitge Hospital (no. PR178/11).

## Availability of data and materials

We provide free access to the co-methylation network data at http://maplab.cat/corre without registration. As for reproducibility, source code is released under the GPL free software license at http://bitbucket.org/imallona/correlations (analysis) and http://bitbucket.org/imallona/corre (shiny app).

Methylation data from the Colonomics project can be downloaded from GEO with accession GSE131013; TCGA data are available at https://portal.gdc.cancer.gov.

## Authors’ contributions

IM, VM and MAP conceived the study. IM developed the method and wrote the software. IM, SA, ADV, VM and MAP analyzed data. IM and MAP wrote the manuscript. All authors read and approved the manuscript.

## Acknowledgements

We thank Iñaki Martinez de Ilarduya and Judith Flo for their excellent technical support and Francisco Chen for contributing to the sequential correlations parallelization. Results presented here are in part based upon data generated by the TCGA Research Network: http://cancergenome.nih.gov/.

## Authors’ information

Izaskun Mallona is currently a SIB Swiss Institute of Bioinformatics and University of Zurich member (Department of Molecular Mechanisms of Disease DMMD, and Institute of Molecular Life Sciences DMLS).

