## Supplementary material for "DNA co-methylation networks outline the structure and remodeling dynamics of colorectal cancer epigenome"

### Contents

1 Supplementary methods 3
1.1 Data 3
1.2 Sequential correlations 3
1.3 Multiple testing adjustment 3
1.4 Network model 4
1.5 Power law fitting 5
1.6 PageRank 5
1.7 Co-methylome and network conservation 5
1.7.1 CpG sharedness visualization 5
1.7.2 Modules preservation analysis by cross-tabulation and Rand index 5
1.7.3 Co-methylation modules conservation visualization 6
1.8 Overlap to partially methylated domains, DMRs and small DMRs 6
1.9 HiC-like visualization 6
1.10 A/B compartments 7
1.11 Purity evaluation and correction 7
1.12 Imprinting 7
1.13 Enrichment tests to Molecular signatures database (MSigDB) 8
1.14 Motif search and enrichment analysis 8
1.15 Co-methylation browser 9
1.16 Statistical significance 9
2 Supplementary tables 10
3 Supplementary figures 18

### 1 Supplementary methods

#### 1.1 Data

We retrieved level 3 DNA methylation normalized data from The Cancer Genome Atlas as evaluated by the Infinium 450k array using TCGA-Assembler (Zhu et al., 2014). Gene expression data were fetched as RNASeq counts (level 3) from the GDC data portal (htseq gene counts). DNA methylation data included COAD (colon), PRAD (prostate adenocarcinoma), LUAD (lung adenocarcinoma), THCA (thyroid cancer) and BRCA (breast cancer).

Colon tumors and adjacent normal data were retrieved from the Colonomics project (Sole et al., 2014; Cordero et al., 2014; Closa et al., 2014). Briefly, frozen samples from primary adenocarcinomas and adjacent normal mucosa were retrieved from the same subjects at Bellvitge University Hospital (Barcelona, Spain) between January 1996 and December 2000. Gene expression was evaluated by the Affymetrix Human Genome U219 chips (Cordero et al., 2014); DNA methylation by the Infinium Illumina 450k array; and exomes as described in (Sanz-Pamplona et al., 2015).

Batch effects were assessed by visually inspecting the unsupervised clustering of a random subset of the 10% of the Infinium probes as compared to the plate ids (Figure S3).

#### 1.2 Sequential correlations

Sequential Spearman's correlation analysis of the methylation status of pairs of reliable probes available at the Infinium HumanMethylation450K arrays, after filtering out for unreliable sites (Price et al., 2013) and probes with missing data or low variability ($sd < 0.05$). That is, a sequential seek of association between probes no matter the actual distance between them. Significant correlations were stored in a PostgreSQL relational database.

#### 1.3 Multiple testing adjustment

To evaluate the $\rho$ effect sizes significance after multiple testing, a cohort-specific Bonferroni-adjusted cut-off was built dividing the signification threshold $\alpha = 0.05$ by the number of total tests. As asymptotic p-values can be calculated from $\rho$ values using the Fisher' Z transform (Fisher, 1915), we searched for the minimum

$\rho$ value to fulfill the Bonferroni-adjusted threshold (one dimensional optimization).

The Colonomics cohort required $251733^2 = 63,369,503,289$ pairwise Spearman's correlations, and required subsequent probability recalculation addressing the multiple testing problem. The Fischer Z transformation of the correlation coefficient $\rho$ according to the cohort size allows to translate the estimate to a pvalue (Fisher, 1915; Shakhbazov et al., 2016), which can be Bonferroni corrected for multiple testing. For instance, the Colonomics tumor series consists on 90 samples with 251,733 variable CpGs which, accounting for multiple testing, led to an adjusted p-value cut-off of $0.05/(251733^2) = 7.890231e - 13$ which is achieved with $\rho \geq 0.8338085$ (see table S1).

##### 1.4 Network model

Given the high number of Spearman correlations run (i.e. $251733^2 = 63,369,503,289$ for Colonomics tumor), we favoured setting astringent correlations cut-offs prior to building the weighted networks. We note that unfiltered weighted networks of smaller datasets have been successfully used in cancer settings, specially for co-expression (Langfelder and Horvath, 2008).

Hence, top-scoring correlations (effect size $\rho \geq 0.8$) were modelled as a graph (network) whose nodes (vertex) are the loci and the edges, the correlations between them. This approach adds data mining flexibility due two reasons: first, the probes have biological meaningful attributes (such as the genomic compartment of the probe, or further associations with, for instance, clinical data or expression); second, the association between probes (the edges, the correlations) reflect the nodes' attributes plus the interactions between them.

We took advantage of the network modularity to detect highly connected compartments (named modules or communities) of the highest correlating probes ($\rho \geq 0.8$) using the fast-greedy algorithm (Clauset et al., 2004). Briefly, the fast greedy algorithm is a bottom-up hierarchical clustering approach which starts with solitary nodes and merges them into the same community if, when doing so, the modularity increase is locally optimal (that is, yields the highest value). Network analysis was performed with igraph_1.0.1 on R 3.2.0 (Csardi and Nepusz, 2006). Plots were rendered with Cytoscape v3.2.1 (Shannon et al., 2003).

#### 1.5 Power law fitting

Degree distributions were inspected using Complementary Cumulative Distribution Functions (CCDF) (Clauset et al., 2009) and statistically evaluated using the R packages fitdistrplus v1.0-9 (Delignette-Muller and Dutang, 2015) and poweRlaw_0.60.3 (Gillespie, 2014).

#### 1.6 PageRank

Google's PageRank was calculated using the igraph (Csardi and Nepusz, 2006) prpack implementation. Damping factor was set in 0.85 (default).

#### 1.7 Co-methylome and network conservation

##### 1.7.1 CpG sharedness visualization

To depict the shared CpGs across modules, we used circular table views rendered by Circos v0.68-pre1 (Krzywinski et al., 2009); and Sankey diagrams from the googleVis R package v0.6.2 (Gesmann and de Castillo, 2011).

##### 1.7.2 Modules preservation analysis by cross-tabulation and Rand index

As each cohort was subjected independently to the correlation analysis and the network modeling, we checked first whether the graphs shared an overall community structure; and second, community by community, which were the closest matches between independent cohorts.

To compare the communities structure at different cohorts, we retrieved the whole co-methylome probes for each cohort as well the community they belong to and then computed partition similarities according to the adjusted Rand index. The adjusted Rand statistic is a distance measure easily interpreted as a probability, being zero when the congruence is expected by chance and one when the matching is perfect (Hubert and Arabie, 1985).

We tested module preservation across cohorts using a cross-tabulation approach (Langfelder et al., 2011). Briefly, the test is based on one-sided Fisher exact tests on whether the CpGs present at two modules do show an overlap greater than what expected by chance. That is, for each cohort, we took the Colonomics tumor partitioning into modules as the standard and compared them to as many modules as present at the evaluated cohort (i.e. Colonomics normal, TCGA normal and TCGA tumor).

##### 1.7.3 Co-methylation modules conservation visualization

To draw the co-methylation modules conservation across datasets (i.e. figure S23) we retrieved the CpGs belonging to each Colonomics tumor module and plotted the network of significant correlations between its members. These networks are not expected to be complete graphs (i.e. linked to saturation) even at the Colonomics tumor dataset, as not necessarily all its CpGs must co-methylate with each other. Nonetheless, given the fact that the module was built on these data, their expected connectivity at Colonomics tumor is very dense. Therefore, we represented side by side the networks for the yardstick (Colonomics tumor), the Colonomics normal and the validation series for normal and tumor at TCGA; some CpGs are not available for some cohorts due to quality filters.

#### 1.8 Overlap to partially methylated domains, DMRs and small DMRs

To look for overlap to IMR90 partially methylated domains (PMDs) we retrieved the regions coordinates from the Lister and Pelizzola (Lister et al., 2009) supplementary data, discarded sex chromosomes events and evaluated their overlap to the co-methylating CpG coordinates using bedtools.

Hansen's DMRs and small DMRs were retrieved from supplemental tables 3 and 7 from Hansen et al. (2011), respectively.

To statistically test their significance, we accounted for the Infinium 450k CpG distribution bias, we run 10,000 permutation tests using the regioneR package (Gel et al., 2015) using the CpGs found to be variable enough (DNA methylation beta value of $SD \geq 0.05$ at the Colonomics tumor cohort) as background (table S7).

#### 1.9 HiC-like visualization

To evaluate the correlation matrix as in HiC analysis, we segmented each chromosome in bins of 1 Mbp and counted the number of significant ($\rho \geq 0.8$) co-methylations between each pair of bins placed at the same chromosome, without any normalization for distance effects. Raw count matrices were plotted with

HiCPlotter v0.7.3 (Akdemir and Chin, 2015), including insulation scores calculation.

##### 1.10 A/B compartments

For TCGA tumor data, Fortin's colon cancer genome segmentation in A/B compartments were retrieved from (Fortin and Hansen, 2015). Segmentation of Colonomics' datasets were carried out using the `compartments()` function at minfi v1.22.1 (Aryee et al., 2014) under R v3.4.0 with the following parameters: 100,000 bp resolution, Spearman correlation method and open sea probes.

##### 1.11 Purity evaluation and correction

To evaluate the influence of tumor purity (e.g. the extent of normal tissue contamination at the tumoral samples) at the DNA co-methylation network we compared the DNA correlation coefficients distribution of raw DNA methylation values with purity-corrected readouts. Briefly, such method evaluates the degree of tissue mixture and corrects for the non-tumor contribution using the observed DNA methylation from matched normal tissue, as well to the malignancy type (colon, COAD code at TCGA). Purity correction was conducted with InfiniumPurify v1.3.1 indicating both tumor and matched normal datasets and the tumor type (Zheng et al., 2017) to increase sensitivity.

When applied to the TCGA dataset, tumor purities estimations were consistent with immunohistological (IHC) and consensus purity measure from (Aran et al., 2015). Colonomics and TCGA purities are depicted at Figure S18.

##### 1.12 Imprinting

A list with 249 imprinted genes in human was download on June the 23rd 2017 from `http://www.geneimprint.com`. Genes annotated as 'Not imprinted' were removed, leaving 234 gene symbols with known or predicted imprinting status. Their genomic locations were retrieved from the UCSC hg19.knownGene and kgXref MySQL tables, aggregating the full span of their transcripts when necessary.

Probes with at least a co-methylation over $\rho \geq 0.8$ (Colonomics tumor dataset) were sorted according to the number of different co-methylation partners (degree). Genomic locations of the top percentile (highest degree) were searched for overlap against the imprinted genes' using bedtools intersect (Quinlan and Hall,

2010).

##### 1.13 Enrichment tests to Molecular signatures database (MSigDB)

To check for functional enrichment of a set of CpGs, we assigned the closest gene to each CpG and compared the gene list to the molecular signatures database MSigDB version 3.0. MSigDB contains curated gene sets that are significantly associated with particular traits, including chromosome physical location, pathways, transcription factor motifs, co-expression modules and Gene Ontology terms (Liberzon et al., 2011). Enrichment was evaluated by cumulative hypergeometric tests corrected for multiple testing by FDR using Homer (Heinz et al., 2010) .

To check for enrichment in chromatin states, we retrieved chromatin segmentations for human stem cells (hESC) from the UCSC MySQL database (hg19.wgEncodeAwgSegmentationChromhmmH1hesc table), which was simplified into the recommended candidate annotations (Tss or TssF, Active Promoter; PromF, Promoter Flanking; PromP, Inactive Promoter; Enh or EnhF, Candidate Strong enhancer; EnhWF, EnhW, DNaseU, DNaseD or FaireW, Candidate Weak enhancer/DNase; CtrcfO or Ctcf, Distal CTCF/Candidate Insulator; Gen5', Elon, ElonW, Gen3', Pol2 or H4K20, Transcription associated; Low, Low activity proximal to active states; ReprD, Repr or ReprW, Polycomb repressed; and Quies or Art, Heterochromatin/Repetitive/Copy Number Variation). Chromatin colors were assigned to CpGs by simple overlap using bedtools (Quinlan and Hall, 2010).

Additional functional enrichment analyses (for Figure S30) were performed using Metascape (Zhou et al., 2019), WebGestalt (Wang et al., 2013), and GOrilla (Eden et al., 2009) web tools with default parameters.

##### 1.14 Motif search and enrichment analysis

Motif search was conducted using Homer (Heinz et al., 2010). Modules are defined upon co-methylated loci having its members placed long distance apart (different chromosomes or over 1 Mbp), but, still, they can present close-by members, e.g. multiple CpGs of the same promoter. Therefore, we evaluated motif enrichment in 200 bp regions centered in each CpG member. For close CpGs, we standardized the regions subjected to motif search in order to reduce the chance of false positives due to counting the same genomic feature multiple times. To do so, we merged overlapping regions into wider intervals and sliced them into 200 bp-long non overlapping intervals (flanks were slopped to produce equally sized regions, when necessary). The 200bp regions were taken as target sequences for the analysis. To look for motif enrichment, we compared the number of hits against a background with matching GC% content to each module sequence set.

##### 1.15 Co-methylation browser

To disseminate results, we developed corre, an R/shiny app freely accessible at http://maplab.cat/corre with any modern Web browser, including mobile devices, without the install of any dedicated software nor registration. The application queries a PostgreSQL database and renders the resulting data, being therefore the calculations performed server-side. The source code of the browser is released under the GPL v2 terms at https://bitbucket.org/imallona/corre.

Dependencies include shiny v0.12.2, RCircos v1.2.0 (Zhang et al., 2013), RPostgreSQL v0.6-2 (DBI v0.3.1) (Conway et al., 2016) and googleVis v0.6.2 (Gesmann and de Castillo, 2011).

##### 1.16 Statistical significance

Two tailed-tests, significance cut-offs of $\alpha = 0.05$ and multiple testing correction by Benjamini and Hochberg were used for hypothesis testing, unless stated otherwise.

### 2 Supplementary tables

Table S1: Statistical cut-offs for simultaneous correlations (multiple testing). `Schema`, the dataset. `N`, the number of samples. `num variable probes`, the number of probes with enough variability and therefore included into the sequential correlations analysis. `adjusted p-value threshold`, the Bonferroni-corrected threshold with as defined by $0.05/(numvariableprobes)^2$. `rho threshold`, the correlation coefficient that fullfills the adjusted p-value threshold as calculated by Fisher Z-transformation optimization (Fisher, 1915; Shakhbazov et al., 2016).

| schema | N | num variable probes | adjusted p-value threshold | $\|\rho\|$ threshold |
|---|---|---|---|---|
| colonomics tumor | 90 | 251733 | 7.890231e-13 | 0.8338085 |
| colonomics normal | 90 | 99346 | 5.066047e-12 | 0.8338085 |
| tcga coad tumor | 256 | 245981 | 8.263554e-13 | 0.6180343 |
| tcga coad normal | 38 | 122398 | 3.337501e-12 | 0.9578317 |

Table S2: Network characteristics. The networks are built using only positive correlations with $\rho \geq 0.8$. `Graph density`, the ratio number of edges / number of possible edges. `diameter`, the longest geodesic. `Mean degree`, the average number of connections by vertex. `Ecount` and `vcount`, the number of edges and vertices, respectively. `Transitivity`, the clustering coefficient, being the probability of the adjacent vertices of a vertex to be connected. `Modularity`, score.

| network | graph density | diameter | mean degree | vcount | ecount | transitivity | modularity |
|---|---|---|---|---|---|---|---|
| colonomics tumor | 0.013 | 25.64 | 835.34 | 63130 | 26367429 | 0.60 | 0.47 |
| colonomics normal | 0.007 | 20.27 | 316.42 | 46885 | 7417715 | 0.54 | 0.62 |
| tcga coad tumor | 0.012 | 14.86 | 450.02 | 37277 | 8387612 | 0.61 | 0.41 |
| tcga coad normal | 0.012 | 15.67 | 936.60 | 76024 | 35601852 | 0.59 | 0.53 |

Table S3: Colonomics tumor modules network description.

| module id | num nodes | num edges | diameter | mean degree |
|---|---|---|---|---|
| 1 | 15,259 | 9,119,877 | 7.35 | 1195.34 |
| 2 | 18,727 | 16,683,557 | 5.74 | 1781.77 |
| 3 | 5,099 | 344,984 | 17.30 | 135.31 |
| 4 | 6,750 | 99,054 | 26.47 | 29.35 |
| 5 | 385 | 1,050 | 22.26 | 5.45 |
| 8 | 168 | 395 | 6.52 | 4.70 |
| 12 | 31 | 118 | 4.09 | 7.61 |
| 13 | 15 | 32 | 4.36 | 4.27 |
| 26 | 11 | 23 | 4.17 | 4.18 |
| 32 | 12 | 36 | 3.44 | 6.00 |
| 41 | 12 | 25 | 3.53 | 4.17 |
| 46 | 14 | 47 | 3.34 | 6.71 |
| 47 | 25 | 35 | 8.31 | 2.80 |
| 64 | 19 | 45 | 4.96 | 4.74 |
| 71 | 13 | 16 | 4.17 | 2.46 |
| 83 | 24 | 93 | 4.19 | 7.75 |
| 120 | 18 | 37 | 4.06 | 4.11 |
| 152 | 28 | 129 | 4.25 | 9.21 |
| 174 | 16 | 39 | 3.44 | 4.88 |
| 184 | 10 | 18 | 2.58 | 3.60 |
| 190 | 13 | 30 | 4.20 | 4.62 |
| 192 | 36 | 56 | 6.69 | 3.11 |
| 198 | 14 | 18 | 4.32 | 2.57 |
| 218 | 62 | 374 | 5.81 | 12.06 |
| 270 | 16 | 37 | 4.34 | 4.62 |
| 276 | 10 | 21 | 3.34 | 4.20 |
| 322 | 16 | 40 | 3.36 | 5.00 |
| 327 | 11 | 22 | 3.32 | 4.00 |
| 598 | 37 | 115 | 5.19 | 6.21 |
| 2352 | 32 | 129 | 3.44 | 8.06 |
| 2992 | 11 | 55 | 0.98 | 10.00 |
| 3237 | 12 | 66 | 0.99 | 11.00 |

Table S4: Module members (CpGs) genomic locations.

| module_id | num_probes | probes_location |
|---|---|---|
| 1 | 15259 | chr1:1338, chr10:838, chr11:908, chr12:779, chr13:433, chr14:524, chr15:473, chr16:532, chr17:719, chr18:316, chr19:768, chr2:1090, chr20:311, chr21:160, chr22:148, chr3:813, chr4:858, chr5:1065, chr6:1217, chr7:975, chr8:746, chr9:248 |
| 2 | 18727 | chr1:2080, chr10:938, chr11:1387, chr12:982, chr13:377, chr14:667, chr15:574, chr16:891, chr17:1119, chr18:163, chr19:842, chr2:1371, chr20:340, chr21:187, chr22:349, chr3:1007, chr4:815, chr5:1005, chr6:1311, chr7:1132, chr8:825, chr9:365 |
| 3 | 5099 | chr1:559, chr10:303, chr11:361, chr12:227, chr13:93, chr14:163, chr15:216, chr16:169, chr17:326, chr18:96, chr19:192, chr2:425, chr20:128, chr21:59, chr22:129, chr3:246, chr4:280, chr5:213, chr6:338, chr7:257, chr8:223, chr9:96 |
| 4 | 6750 | chr1:699, chr10:331, chr11:581, chr12:298, chr13:149, chr14:177, chr15:216, chr16:276, chr17:334, chr18:25, chr19:274, chr2:592, chr20:231, chr21:36, chr22:142, chr3:407, chr4:245, chr5:332, chr6:559, chr7:500, chr8:232, chr9:114 |
| 5 | 385 | chr1:14, chr10:23, chr11:54, chr12:7, chr13:6, chr14:15, chr15:22, chr16:17, chr17:8, chr18:19, chr19:17, chr2:10, chr20:34, chr22:2, chr3:57, chr4:25, chr5:11, chr6:12, chr7:23, chr8:9 |
| 8 | 168 | chr1:10, chr10:16, chr11:9, chr12:13, chr14:6, chr15:8, chr16:2, chr17:5, chr18:4, chr2:16, chr20:1, chr3:10, chr4:10, chr5:24, chr6:4, chr7:14, chr8:16 |
| 12 | 31 | chr1:2, chr10:2, chr12:1, chr14:2, chr16:4, chr17:1, chr19:2, chr2:7, chr20:1, chr4:2, chr5:1, chr6:3, chr7:2, chr9:1 |
| 13 | 15 | chr1:15 |
| 26 | 11 | chr14:8, chr2:1, chr9:2 |
| 32 | 12 | chr13:12 |
| 41 | 12 | chr15:5, chr20:7 |
| 46 | 14 | chr19:11, chr2:1, chr7:1, chr8:1 |
| 47 | 25 | chr1:3, chr10:8, chr15:5, chr19:1, chr2:3, chr6:5 |
| 64 | 19 | chr15:3, chr17:1, chr5:15 |
| 71 | 13 | chr10:1, chr11:1, chr17:4, chr2:1, chr22:2, chr6:1, chr9:3 |
| 83 | 24 | chr1:11, chr12:13 |
| 120 | 18 | chr1:3, chr11:1, chr12:2, chr13:1, chr19:3, chr2:2, chr6:5, chr7:1 |
| 152 | 28 | chr10:4, chr14:1, chr18:19, chr22:4 |
| 174 | 16 | chr10:11, chr4:5 |
| 184 | 10 | chr10:5, chr11:5 |
| 190 | 13 | chr12:1, chr21:4, chr3:8 |
| 192 | 36 | chr1:6, chr16:3, chr17:12, chr19:1, chr2:11, chr9:3 |
| 198 | 14 | chr1:8, chr11:1, chr17:2, chr8:3 |
| 218 | 62 | chr20:62 |
| 270 | 16 | chr11:15, chr5:1 |
| 276 | 10 | chr20:1, chr4:9 |
| 322 | 16 | chr7:16 |
| 327 | 11 | chr12:10, chr5:1 |
| 598 | 37 | chr10:9, chr11:12, chr15:1, chr17:4, chr19:4, chr2:4, chr22:3 |
| 2352 | 32 | chr12:7, chr18:21, chr21:4 |
| 2992 | 11 | chr11:10, chr19:1 |
| 3237 | 12 | chr14:10, chr16:1, chr9:1 |

Table S5: Modules top MSigDB annotation.

| module id | p-value | term | go_tree | goi_id | term_n | term_target_n | n | target_n |
|---|---|---|---|---|---|---|---|---|
| 1 | 0.00 | BENPORATH_ES_WITH_H3K27ME3 | MSigDB lists | BENPORATH_ES_WITH_H3K27ME3 | 1066 | 504 | 18039 | 3411 |
| 2 | 0.00 | colon | COSMIC cancer mutations | colon | 17647 | 6278 | 18488 | 6399 |
| 3 | 0.00 | MEISSNER_BRAIN_HCP_WITH_H3K4ME | MSigDB lists | MEISSNER_BRAIN_HCP_WITH_H3K4ME3_AND_H3K27ME3 | 1044 | 273 | 18039 | 2089 |
| 4 | 0.00 | thyroid | COSMIC cancer mutations | thyroid | 8406 | 1626 | 18488 | 2955 |
| 5 | 0.00 | BENPORATH_SUZ12_TARGETS | MSigDB lists | BENPORATH_SUZ12_TARGETS | 989 | 25 | 18039 | 65 |
| 8 | 0.00 | PILON_KLF1_TARGETS_DN | MSigDB lists | PILON_KLF1_TARGETS_DN | 1912 | 37 | 18039 | 129 |
| 12 | 0.00 | MUELLER_PLURINET | MSigDB lists | MUELLER_PLURINET | 295 | 6 | 18039 | 26 |
| 13 | 0.00 | embryonic hindlimb morphogenes | biological process | GO:0035116 | 31 | 1 | 16088 | 1 |
| 26 | 0.00 | RELA_DN.V1_UP | MSigDB lists | RELA_DN.V1_UP | 144 | 2 | 18039 | 3 |
| 32 | 0.00 | human chr13q13.3 | chromosome location | human chr13q13.3 | 32 | 2 | 47739 | 3 |
| 41 | 0.00 | SKOR1 | interpro domains | IPR028762 | 1 | 1 | 18037 | 2 |
| 46 | 0.00 | NIKOLSKY_BREAST_CANCER_19Q13.1 | MSigDB lists | NIKOLSKY_BREAST_CANCER_19Q13.1_AMPLICON | 21 | 2 | 18039 | 4 |
| 47 | 0.00 | MEISSNER_BRAIN_HCP_WITH_H3K4ME | MSigDB lists | MEISSNER_BRAIN_HCP_WITH_H3K4ME3_AND_H3K27ME3 | 1044 | 5 | 18039 | 7 |
| 64 | 0.00 | human chr5q23-q31 | chromosome location | human chr5q23-q31 | 2 | 1 | 47739 | 3 |
| 71 | 0.00 | labyrinthine layer blood vesse | biological process | GO:0060716 | 20 | 2 | 16088 | 8 |
| 83 | 0.00 | GSE17721_CTRL_VS_CPG_0.5H_BMDM | MSigDB lists | GSE17721_CTRL_VS_CPG_0.5H_BMDM_DN | 188 | 2 | 18039 | 2 |
| 120 | 0.00 | chr6p21 | MSigDB lists | chr6p21 | 299 | 5 | 18039 | 15 |
| 152 | 0.00 | cochlea development | biological process | GO:0090102 | 35 | 2 | 16088 | 4 |
| 174 | 0.00 | human chr10p12.1 | chromosome location | human chr10p12.1 | 35 | 2 | 47739 | 4 |
| 184 | 0.00 | FJX1/FJ | interpro domains | IPR024868 | 1 | 1 | 18037 | 1 |
| 190 | 0.00 | Pept_M12B_ADAM-TS1 | interpro domains | IPR013274 | 1 | 1 | 18037 | 3 |
| 192 | 0.00 | human chr2q11.2-q12.1 | chromosome location | human chr2q11.2-q12.1 | 1 | 1 | 47739 | 7 |
| 198 | 0.00 | TLR5 | interpro domains | IPR027176 | 1 | 1 | 18037 | 3 |
| 218 | 0.00 | chr20q13 | MSigDB lists | chr20q13 | 157 | 2 | 18039 | 2 |
| 270 | 0.00 | Tet-R_TetA/multi-R_MdtG | interpro domains | IPR001958 | 5 | 1 | 18037 | 1 |
| 276 | 0.00 | BMP3/GDF10 | interpro domains | IPR017197 | 2 | 1 | 18037 | 2 |
| 322 | 0.00 | human chr7p12.2 | chromosome location | human chr7p12.2 | 8 | 1 | 47739 | 2 |
| 327 | 0.00 | KANG_IMMORTALIZED_BY_TERT_DN | MSigDB lists | KANG_IMMORTALIZED_BY_TERT_DN | 99 | 2 | 18039 | 2 |
| 598 | 0.00 | BENPORATH_PRC2_TARGETS | MSigDB lists | BENPORATH_PRC2_TARGETS | 619 | 6 | 18039 | 7 |
| 2352 | 0.00 | GAL1_rcpt | interpro domains | IPR003906 | 1 | 1 | 18037 | 3 |
| 2992 | 0.00 | SNX5/SNX6/SNX32 | interpro domains | IPR014637 | 3 | 1 | 18037 | 2 |
| 3237 | 0.00 | GSE-like | interpro domains | IPR022207 | 1 | 1 | 18037 | 2 |

Table S6: Known motif enrichment analysis (top hit for each module).

| Module id | Motif Name | Consensus | P-value | Log P-value | q-value | # Targets | % Targets | # Backgrounds | % Backgrounds |
|---|---|---|---|---|---|---|---|---|---|
| 1 | EWS:ERG-fusion(ETS)/CADO_ES1-EWS:ERG-ChIP-Seq(SRA014231) | ATTTCCTGTN | 1e-57 | -1.314e+02 | 0.0000 | 1034.0 | 8.29% | 1764.6 | 4.91% |
| 2 | Fra1(bZIP)/BT549-Fra1-ChIP-Seq(GSE46166) | NNATGASTCATH | 1e-138 | -3.179e+02 | 0.0000 | 1239.0 | 8.01% | 1253.9 | 3.66% |
| 3 | CEBP(bZIP)/ThioMac-CEBPb-ChIP-Seq(GSE21512) | ATTGCGCAAC | 1e-33 | -7.625e+01 | 0.0000 | 279.0 | 6.62% | 1329.0 | 2.97% |
| 4 | BATF(bZIP)/Th17-BATF-ChIP-Seq(GSE39756) | DATGASTCAT | 1e-20 | -4.706e+01 | 0.0000 | 400.0 | 6.84% | 1820.5 | 4.17% |
| 5 | Pax7(Paired,Homeobox)/Myoblast-Pax7-ChIP-Seq(GSE25064) | TAATCAATTA | 1e-4 | -9.352e+00 | 0.0277 | 4.0 | 1.58% | 42.6 | 0.09% |
| 8 | Phox2a(Homeobox)/Neuron-Phox2a-ChIP-Seq(GSE31456) | YTAATYNRATTA | 1e-2 | -5.394e+00 | 1.0000 | 21.0 | 14.69% | 3751.0 | 7.93% |
| 12 | NRF1(NRF)/MCF7-NRF1-ChIP-Seq(Unpublished) | CTGCGCATGCGC | 1e-2 | -5.577e+00 | 1.0000 | 7.0 | 22.58% | 2677.9 | 6.69% |
| 13 | Gata4(Zf)/Heart-Gata4-ChIP-Seq(GSE35151) | NBWGATAAGR | 1e-3 | -8.132e+00 | 0.0938 | 4.0 | 40.00% | 1916.0 | 3.60% |
| 26 | Tcf21(bHLH)/ArterySmoothMuscle-Tcf21-ChIP-Seq(GSE61369) | NAACAGCTGG | 1e-2 | -4.985e+00 | 1.0000 | 4.0 | 40.00% | 3863.2 | 8.38% |
| 32 | BMYB(HTH)/Hela-BMYB-ChIP-Seq(GSE27030) | NHAACBGYYV | 1e-2 | -6.371e+00 | 0.5457 | 4.0 | 66.67% | 14055.8 | 10.81% |
| 41 | CHR(?)/Hela-CellCycle-Expression | SRGTTTCAAA | 1e-2 | -5.232e+00 | 1.0000 | 2.0 | 28.57% | 1891.7 | 1.64% |
| 46 | Oct4:Sox17(POU,Homeobox,HMG)/F9-Sox17-ChIP-Seq(GSE44553) | CCATTGTATGCAAAT | 1e-1 | -3.942e+00 | 1.0000 | 1.0 | 14.29% | 370.9 | 0.28% |
| 47 | GATA3(Zf),DR8/iTreg-Gata3-ChIP-Seq(GSE20898) | AGATSTNDNNDSAGATAASN | 1e-3 | -7.446e+00 | 0.1862 | 2.0 | 8.70% | 71.9 | 0.16% |
| 64 | GRHL2(CP2)/HBE-GRHL2-ChIP-Seq(GSE46194) | AAACYKGTTWDACMRGTTTB | 1e-2 | -6.414e+00 | 0.5224 | 2.0 | 28.57% | 975.4 | 0.90% |
| 71 | CEBP:CEBP(bZIP)/MEF-Chop-ChIP-Seq(GSE35681) | NTNATGCAAYMNNHTGMAAY | 1e-2 | -5.613e+00 | 1.0000 | 2.0 | 16.67% | 403.5 | 0.76% |
| 83 | Lhx2(Homeobox)/HFSC-Lhx2-ChIP-Seq(GSE48068) | TAATTAGN | 1e-4 | -1.068e+01 | 0.0073 | 6.0 | 37.50% | 1683.1 | 3.99% |
| 120 | HIF-1b(HLH)/T47D-HIF1b-ChIP-Seq(GSE59937) | RTACGTGC | 1e-3 | -7.782e+00 | 0.1331 | 8.0 | 44.44% | 5433.4 | 11.34% |
| 152 | Mef2a(MADS)/HL1-Mef2a.biotin-ChIP-Seq(GSE21529) | CYAAAAATAG | 1e-3 | -8.141e+00 | 0.0930 | 4.0 | 28.57% | 968.3 | 2.44% |
| 174 | E2F7(E2F)/Hela-E2F7-ChIP-Seq(GSE32673) | VDTTTCCCGCCA | 1e-2 | -5.297e+00 | 1.0000 | 3.0 | 27.27% | 1557.9 | 3.34% |
| 184 | Mef2c(MADS)/GM12878-Mef2c-ChIP-Seq(GSE32465) | DCYAAAAATAGM | 1e-2 | -6.678e+00 | 0.4012 | 2.0 | 33.33% | 1588.1 | 0.93% |
| 190 | CArG(MADS)/PUER-Srf-ChIP-Seq(Sullivan_et_al.) | CCATATATGGNM | 1e-2 | -4.643e+00 | 1.0000 | 2.0 | 16.67% | 639.7 | 1.26% |
| 192 | E2F(E2F)/Hela-CellCycle-Expression | TTSGCGCGAAAA | 1e-1 | -4.488e+00 | 1.0000 | 3.0 | 11.11% | 829.3 | 1.74% |
| 198 | Nkx2.2(Homeobox)/NPC-Nkx2.2-ChIP-Seq(GSE61673) | BTBRAGTGSN | 1e-2 | -5.948e+00 | 0.8332 | 7.0 | 53.85% | 7905.0 | 16.91% |
| 218 | ZNF143—STAF(Zf)/CUTLL-ZNF143-ChIP-Seq(GSE29600) | ATTTCCCAGVAKSCY | 1e-2 | -5.073e+00 | 1.0000 | 4.0 | 18.18% | 1600.6 | 3.45% |
| 270 | SPDEF(ETS)/VCaP-SPDEF-ChIP-Seq(SRA014231) | ASWTCCTGBT | 1e-1 | -4.121e+00 | 1.0000 | 3.0 | 50.00% | 9200.5 | 10.09% |
| 276 | GLI3(Zf)/Limb-GLI3-ChIP-Chip(GSE11077) | CGTGGGTGGTCC | 1e-2 | -5.718e+00 | 1.0000 | 2.0 | 33.33% | 4356.0 | 1.51% |
| 322 | Mef2b(MADS)/HEK293-Mef2b.V5-ChIP-Seq(GSE67450) | GCTATTTTTGGM | 1e-2 | -5.254e+00 | 1.0000 | 2.0 | 20.00% | 1230.2 | 1.11% |
| 327 | SPDEF(ETS)/VCaP-SPDEF-ChIP-Seq(SRA014231) | ASWTCCTGBT | 1e-3 | -7.771e+00 | 0.1346 | 5.0 | 62.50% | 4661.8 | 9.95% |
| 598 | Ascl1(bHLH)/NeuralTubes-Ascl1-ChIP-Seq(GSE55840) | NNVVCAGCTGBN | 1e-3 | -7.947e+00 | 0.1129 | 13.0 | 52.00% | 9470.2 | 19.92% |
| 2352 | Fra1(bZIP)/BT549-Fra1-ChIP-Seq(GSE46166) | NNATGASTCATH | 1e-2 | -6.582e+00 | 0.4420 | 3.0 | 21.43% | 704.3 | 1.63% |
| 2992 | Egr1(Zf)/K562-Egr1-ChIP-Seq(GSE32465) | TGCGTGGGYG | 1e-1 | -3.961e+00 | 1.0000 | 3.0 | 75.00% | 31932.5 | 17.64% |
| 3237 | GABPA(ETS)/Jurkat-GABPa-ChIP-Seq(GSE17954) | RACCGGAAGT | 1e-1 | -3.472e+00 | 1.0000 | 3.0 | 42.86% | 17902.2 | 10.74% |

Table S7: Modules association to Hansen's known varible regions in colon cancer (Hansen et al., 2011). Data are summarized by their Z score and log 10 pvalue (in brackets) (permutation analysis as depicted in figure S28).

| module | hypo | hyper | boundaryShift | lossOfRegulation | novelMethylation | other |
|---|---|---|---|---|---|---|
| 1 | -143.61 (-4.00e+00) | 139.66 (4.00e+00) | 327.1 (4.00e+00) | 1081.22 (4.00e+00) | -1.4 (-8.49e-01) | 491.34 (4.00e+00) |
| 2 | -29.75 (-4.00e+00) | -1.89 (-1.52e+00) | 54.36 (4.00e+00) | -1.36 (-9.93e-01) | 72.67 (4.00e+00) | 62.31 (4.00e+00) |
| 3 | -71.65 (-4.00e+00) | 0.25 (3.82e-01) | 42 (4.00e+00) | 2.9 (2.03e+00) | 7.89 (4.00e+00) | 25.81 (4.00e+00) |
| 4 | -20.02 (-4.00e+00) | 27.93 (4.00e+00) | 14.81 (4.00e+00) | 24.01 (4.00e+00) | 15.22 (4.00e+00) | 19.56 (4.00e+00) |
| 5 | -28.5 (-4.00e+00) | 11.03 (4.00e+00) | 87.16 (4.00e+00) | 329.01 (4.00e+00) | -0.22 (-2.03e-02) | 166.69 (4.00e+00) |
| 8 | -16.06 (-4.00e+00) | 2.06 (1.25e+00) | -0.36 (-5.48e-02) | 2.31 (8.86e-01) | -0.14 (-8.64e-03) | -0.5 (-1.07e-01) |
| 12 | -8.11 (-4.00e+00) | -0.63 (-1.73e-01) | -0.14 (-9.04e-03) | -0.16 (-1.14e-02) | -0.06 (-1.78e-03) | -0.21 (-1.98e-02) |
| 13 | -5.5 (-4.00e+00) | -0.44 (-8.24e-02) | 72.18 (4.00e+00) | 63.38 (4.00e+00) | -0.04 (-8.26e-04) | -0.14 (-8.86e-03) |
| 26 | -4.83 (-4.00e+00) | -0.37 (-5.98e-02) | 34.23 (4.00e+00) | -0.09 (-3.75e-03) | -0.04 (-7.39e-04) | 64.82 (4.00e+00) |
| 32 | -1.9 (-1.22e+00) | -0.39 (-6.53e-02) | 150.39 (4.00e+00) | -0.1 (-4.28e-03) | -0.03 (-3.91e-04) | -0.13 (-7.09e-03) |
| 41 | -4.95 (-4.00e+00) | -0.4 (-6.73e-02) | -0.09 (-3.62e-03) | -0.1 (-4.10e-03) | -0.04 (-6.95e-04) | 86.72 (4.00e+00) |
| 46 | -5.36 (-4.00e+00) | 4.39 (1.89e+00) | -0.1 (-4.41e-03) | 99.68 (4.00e+00) | -0.04 (-8.26e-04) | 13.51 (3.70e+00) |
| 47 | -7.24 (-4.00e+00) | -0.57 (-1.38e-01) | 36.47 (4.00e+00) | 34.47 (4.00e+00) | -0.06 (-1.61e-03) | 74.96 (4.00e+00) |
| 64 | -6.25 (-4.00e+00) | 5.76 (2.68e+00) | -0.12 (-5.86e-03) | -0.13 (-6.83e-03) | -0.05 (-9.56e-04) | 79.55 (4.00e+00) |
| 71 | -4.08 (-3.52e+00) | -0.4 (-7.03e-02) | -0.1 (-4.10e-03) | -0.1 (-4.58e-03) | -0.05 (-9.13e-04) | 7.45 (1.76e+00) |
| 83 | -7.09 (-4.00e+00) | 1.33 (5.92e-01) | 82.58 (4.00e+00) | -0.13 (-7.71e-03) | -0.06 (-1.35e-03) | 67.71 (4.00e+00) |
| 120 | -5.59 (-4.00e+00) | -0.48 (-9.92e-02) | -0.11 (-5.59e-03) | -0.11 (-5.64e-03) | -0.04 (-7.82e-04) | -0.16 (-1.07e-02) |
| 152 | -7.61 (-4.00e+00) | -0.59 (-1.51e-01) | 36.38 (4.00e+00) | 26.12 (4.00e+00) | -0.06 (-1.39e-03) | 88.46 (4.00e+00) |
| 174 | -5.32 (-4.00e+00) | -0.44 (-8.24e-02) | -0.12 (-5.90e-03) | 17.82 (3.70e+00) | -0.04 (-8.26e-04) | 91.46 (4.00e+00) |
| 184 | -4.52 (-4.00e+00) | -0.35 (-5.33e-02) | 46.9 (4.00e+00) | -0.09 (-3.62e-03) | -0.04 (-6.08e-04) | 25.88 (4.00e+00) |
| 190 | -5.2 (-4.00e+00) | 4.7 (2.00e+00) | 38.42 (4.00e+00) | 75.58 (4.00e+00) | -0.04 (-7.82e-04) | 7.49 (1.76e+00) |
| 192 | -8.69 (-4.00e+00) | -0.67 (-1.93e-01) | 63.02 (4.00e+00) | -0.16 (-1.12e-02) | -0.07 (-2.13e-03) | -0.23 (-2.20e-02) |
| 198 | -1.42 (-8.83e-01) | -0.42 (-7.67e-02) | 28.27 (4.00e+00) | -0.11 (-5.73e-03) | -0.05 (-9.13e-04) | 14 (3.30e+00) |
| 218 | -11.32 (-4.00e+00) | -0.88 (-3.34e-01) | -0.21 (-1.93e-02) | -0.23 (-2.20e-02) | -0.09 (-3.18e-03) | -0.3 (-3.86e-02) |
| 270 | -5.77 (-4.00e+00) | -0.45 (-8.57e-02) | -0.11 (-5.68e-03) | -0.11 (-5.59e-03) | -0.04 (-8.69e-04) | -0.14 (-9.08e-03) |
| 276 | -4.58 (-4.00e+00) | -0.35 (-5.39e-02) | -0.09 (-3.14e-03) | -0.09 (-3.18e-03) | -0.03 (-3.91e-04) | 79.62 (4.00e+00) |
| 322 | -5.77 (-4.00e+00) | -0.45 (-8.69e-02) | -0.1 (-4.32e-03) | 137.97 (4.00e+00) | -0.05 (-1.09e-03) | -0.15 (-1.01e-02) |
| 327 | -4.74 (-4.00e+00) | -0.36 (-5.70e-02) | 10.27 (2.02e+00) | -0.09 (-3.14e-03) | -0.04 (-6.52e-04) | -0.13 (-7.00e-03) |
| 598 | -8.66 (-4.00e+00) | -0.7 (-2.08e-01) | -0.17 (-1.30e-02) | 166.32 (4.00e+00) | -0.07 (-2.05e-03) | 12.21 (4.00e+00) |
| 2352 | -8.2 (-4.00e+00) | -0.63 (-1.72e-01) | -0.15 (-1.05e-02) | 151.74 (4.00e+00) | -0.05 (-1.09e-03) | 22.67 (4.00e+00) |
| 2992 | -4.77 (-3.70e+00) | -0.37 (-5.94e-02) | -0.09 (-3.88e-03) | 101.91 (4.00e+00) | -0.04 (-6.08e-04) | -0.13 (-6.83e-03) |
| 3237 | -4.88 (-4.00e+00) | -0.38 (-6.39e-02) | 95.2 (4.00e+00) | -0.1 (-3.97e-03) | -0.05 (-1.00e-03) | -0.13 (-6.87e-03) |

### 3 Supplementary figures

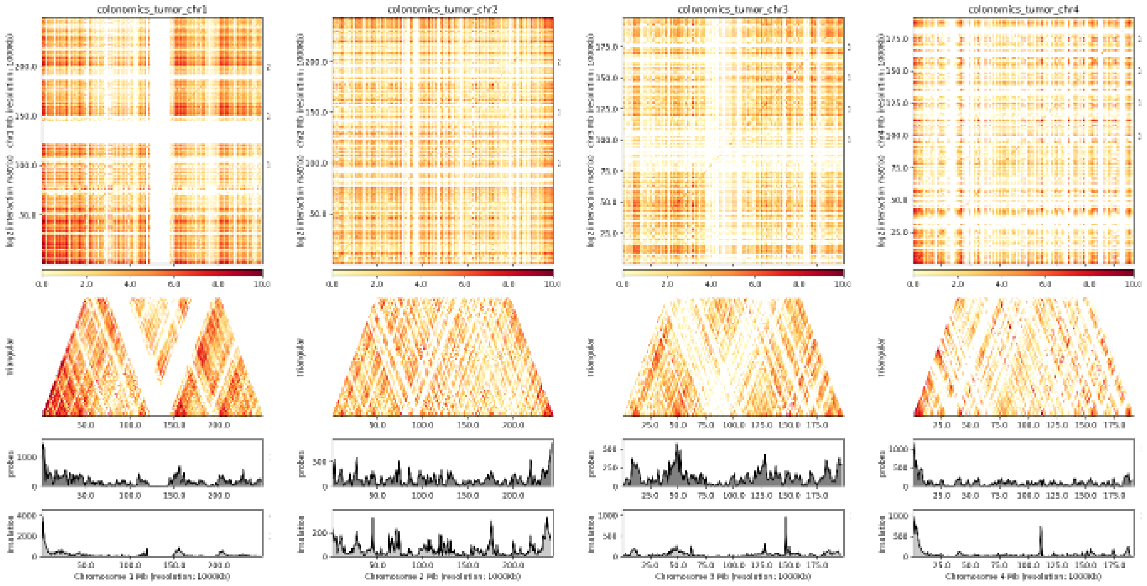

Figure S1: HiC-like visualization. Positive correlations do not decay with distance and their distribution differs from the local probes'. From left to right: chromosomes 1 to 4 binned in 1 Mbp-long segments. Tracks depict a triangular view of the whole matrix, the Infinium 450k probe density and the insulation score from the cis-comethylating probes.

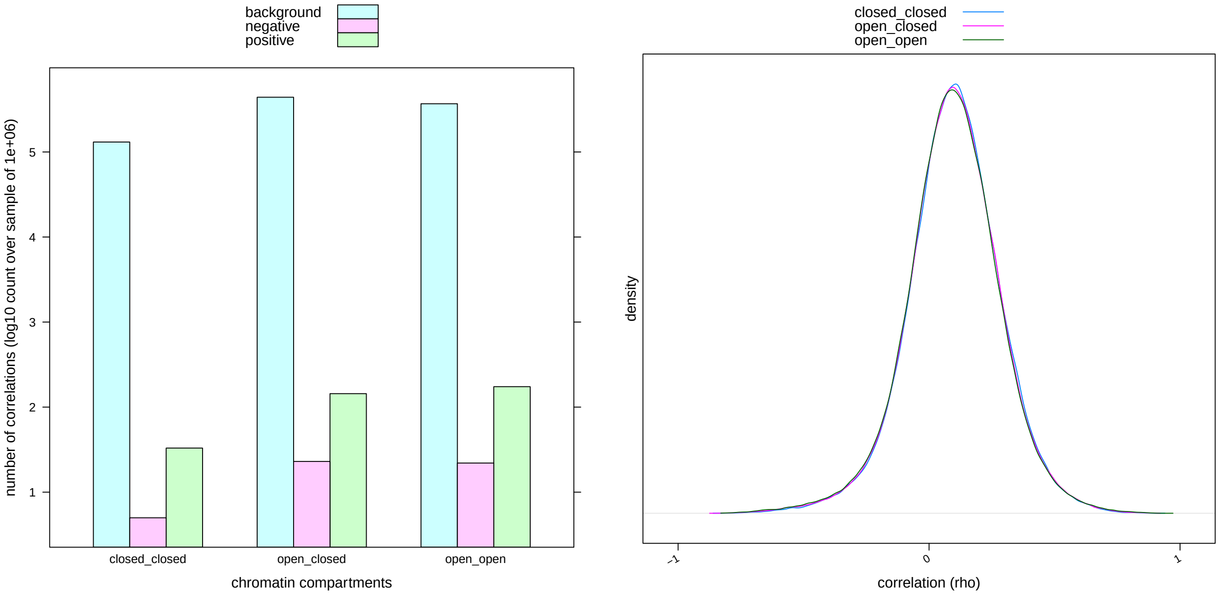

Figure S2: Correlations distribution by A/B compartments. A, correlations with at least one CpG placed within an open compartment show higher number of significant co-methylations (background: a million intra-chromosomal correlations sampled from chromosome 10; negative: those with $\rho < -0.8$; positive: those with $\rho \geq 0.8$). B, the bell-shape distribution does not depend on the open or closed chromatin status of the comethylated-probes. Chromatin compartments were estimated from DNA methylation values according to (Fortin and Hansen, 2015).

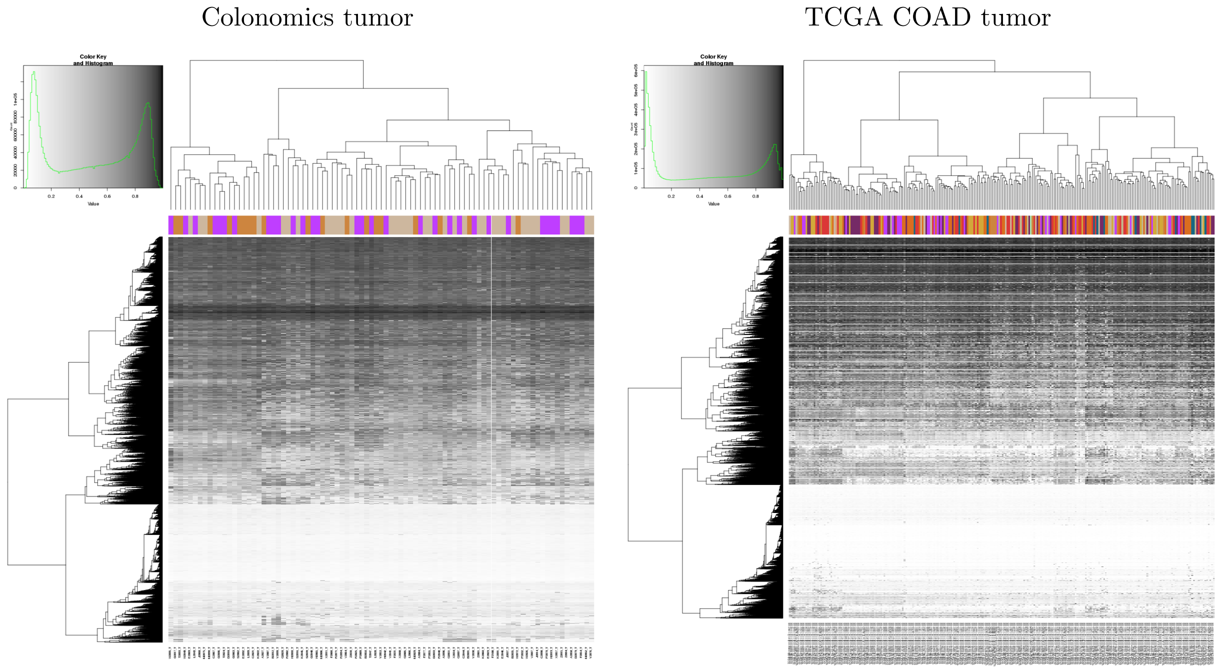

Figure S3: Batch effect assessment. The unsupervised clustering of DNA methylation at the Colonomics cohort (left) nor the TCGA coad (right) does not group samples processed at the same batch (plate id, colored columns). Hyerarchical clustering (Ward method, ward.D2 in base R) of Euclidean distance matrix built upon the methylation status of a random subset composed by the 10% of the Infinium probes.

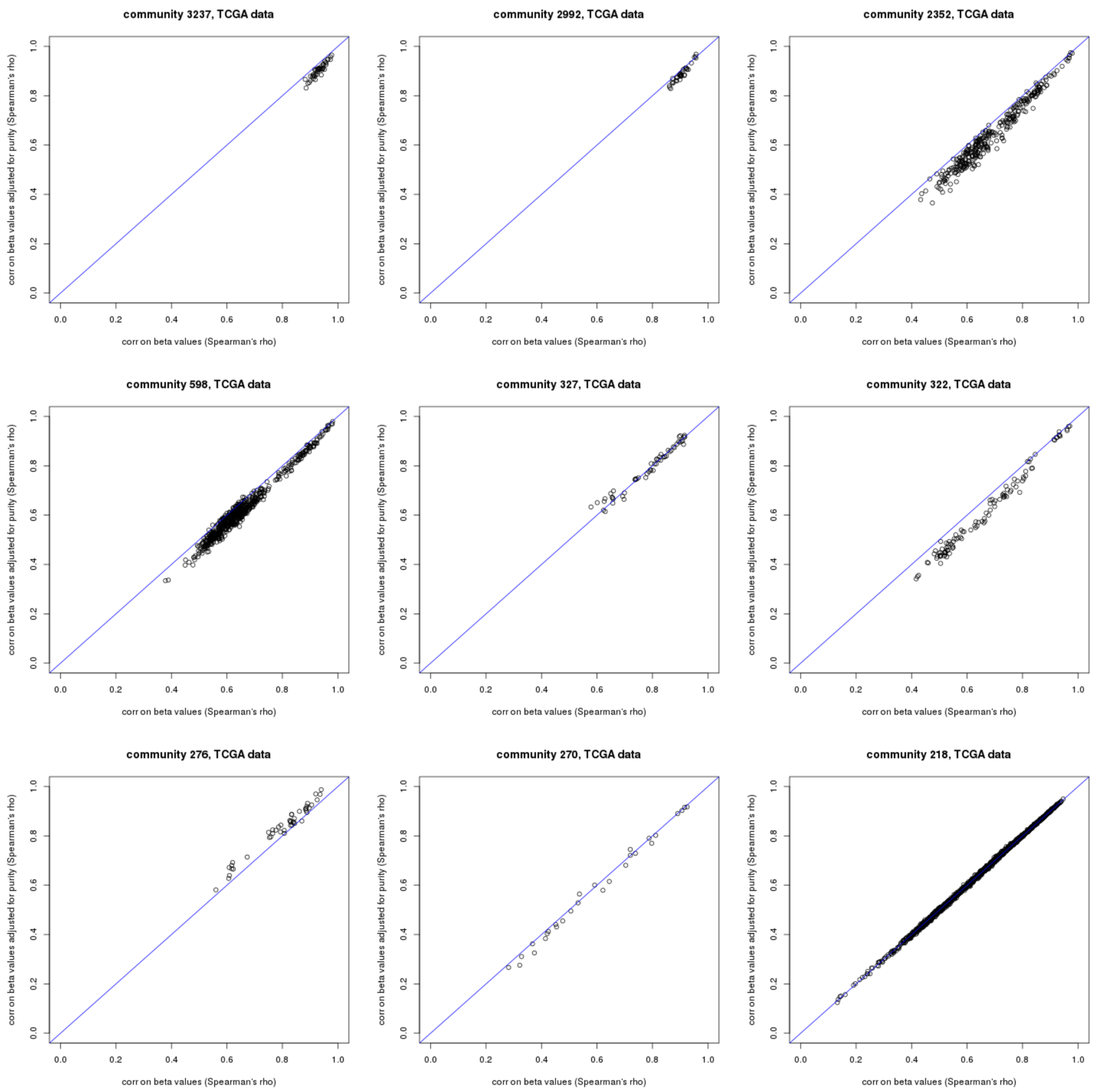

Figure S4: Purity linear model subtraction. Correlation structure is maintained after removal of purity effects by a linear model. For the sake of brevity only nine communities are shown.

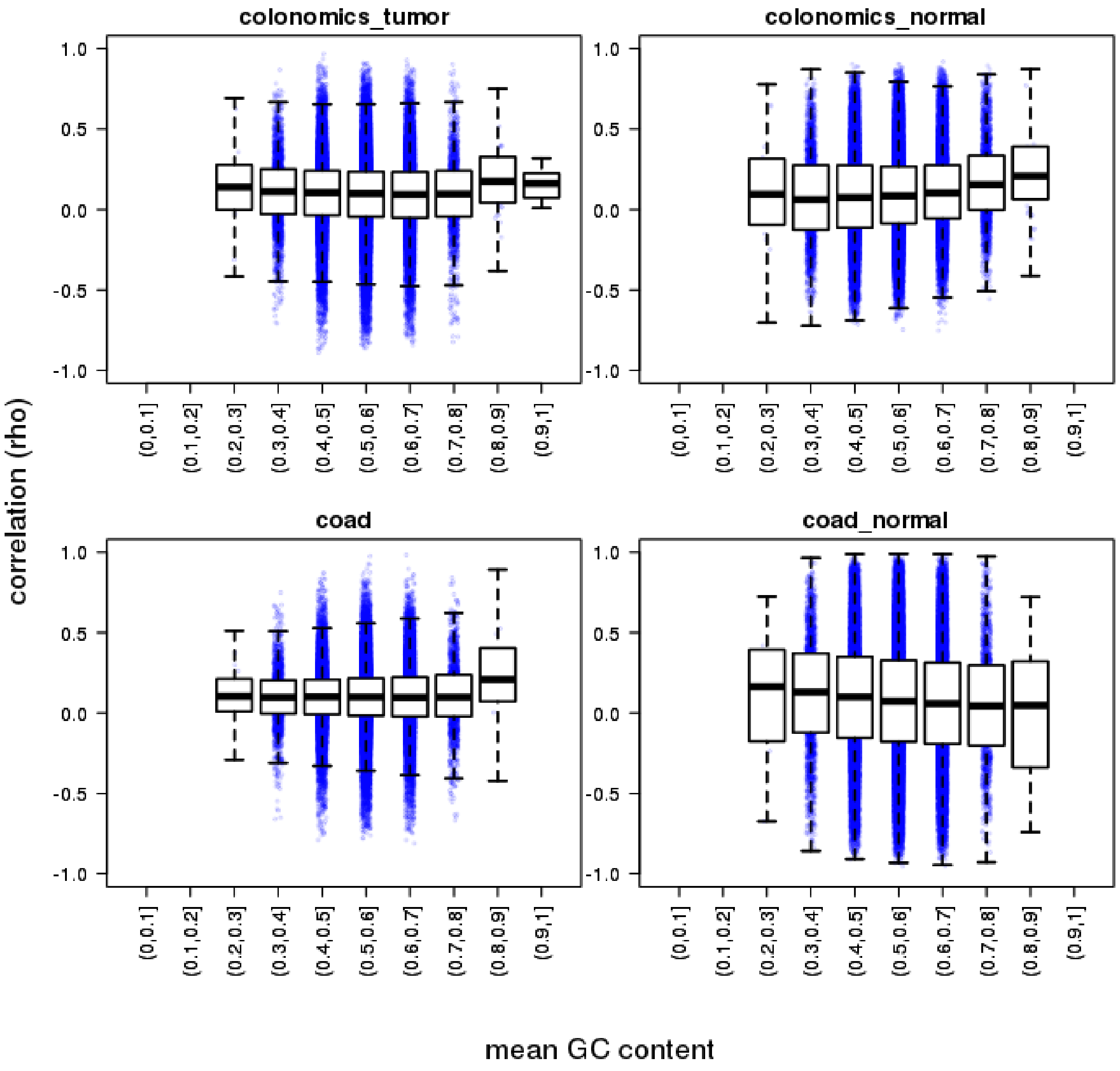

Figure S5: Correlation coefficients stratified by the Infinium probes GC content. Comprehensive unfiltered correlations were calculated for the chr9 vs chr10 and the chr10 against itself. GC values for each probe pair were averaged and stratified. Boxplots depict the full dataset; the stripchart is built upong a random sample of 100,000 probe pairs.

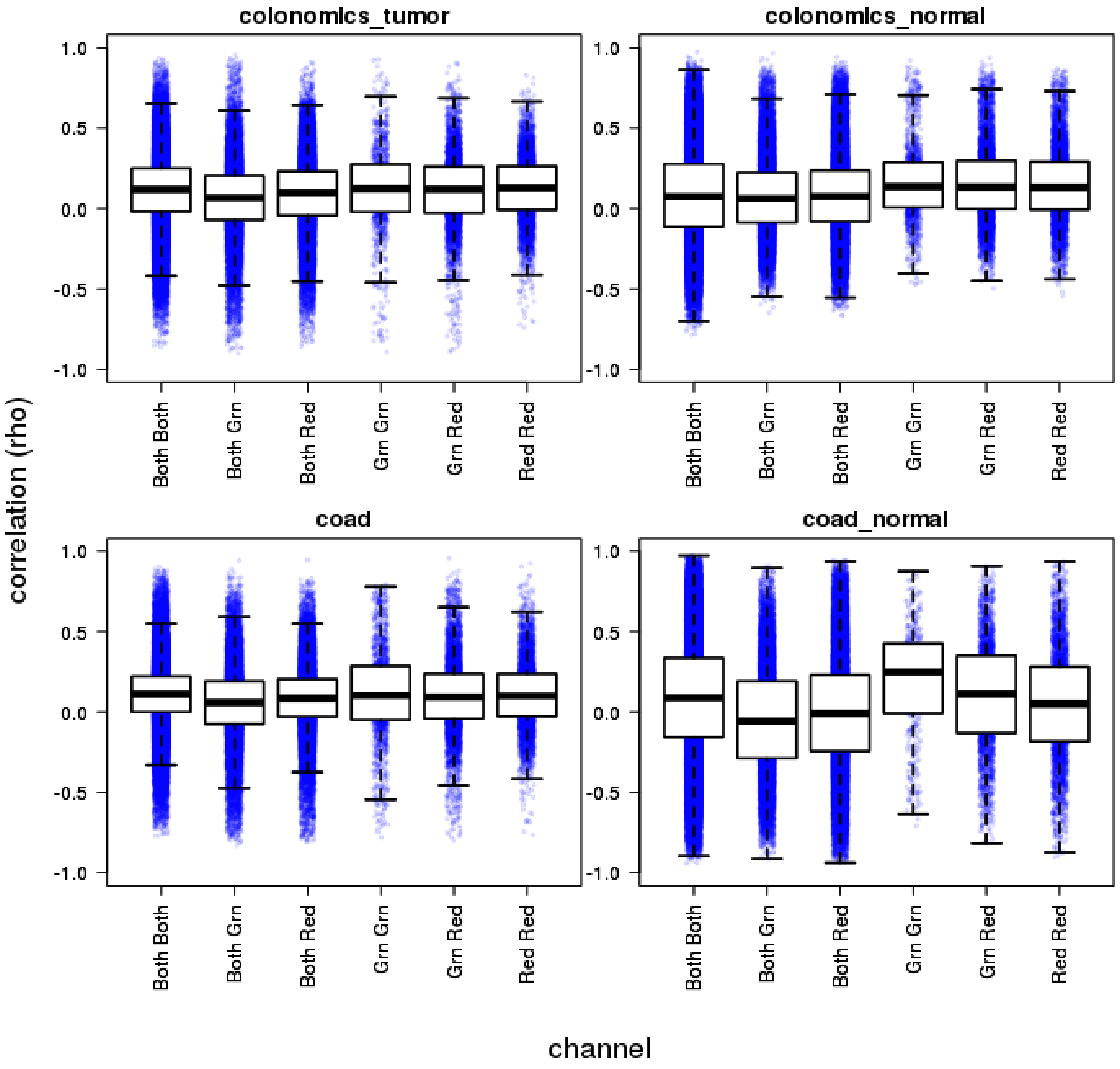

Figure S6: Correlation coefficients stratified by the Infinium probes measuring channel. Comprehensive unfiltered correlations were calculated for the chr9 vs chr10 and the chr10 against itself. Boxplots depict the full dataset; the stripchart is built upong a random sample of 100,000 probe pairs in total.

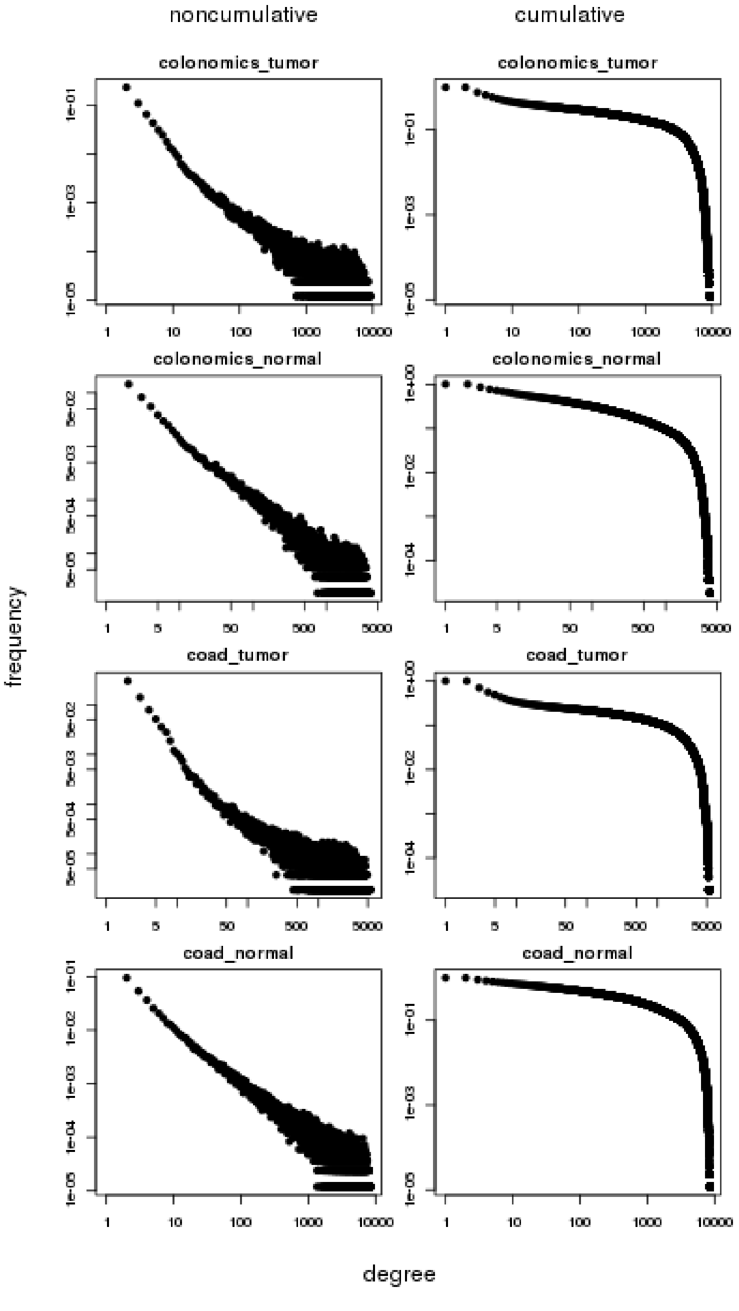

Figure S7: Degree distribution. Log-log plots of the degree distribution (left) and the complementary cumulative distribution function (CCDF; right) for the four datasets. The lack of linearity of the CCDF rules out a power law behaviour along the whole degree distribution.

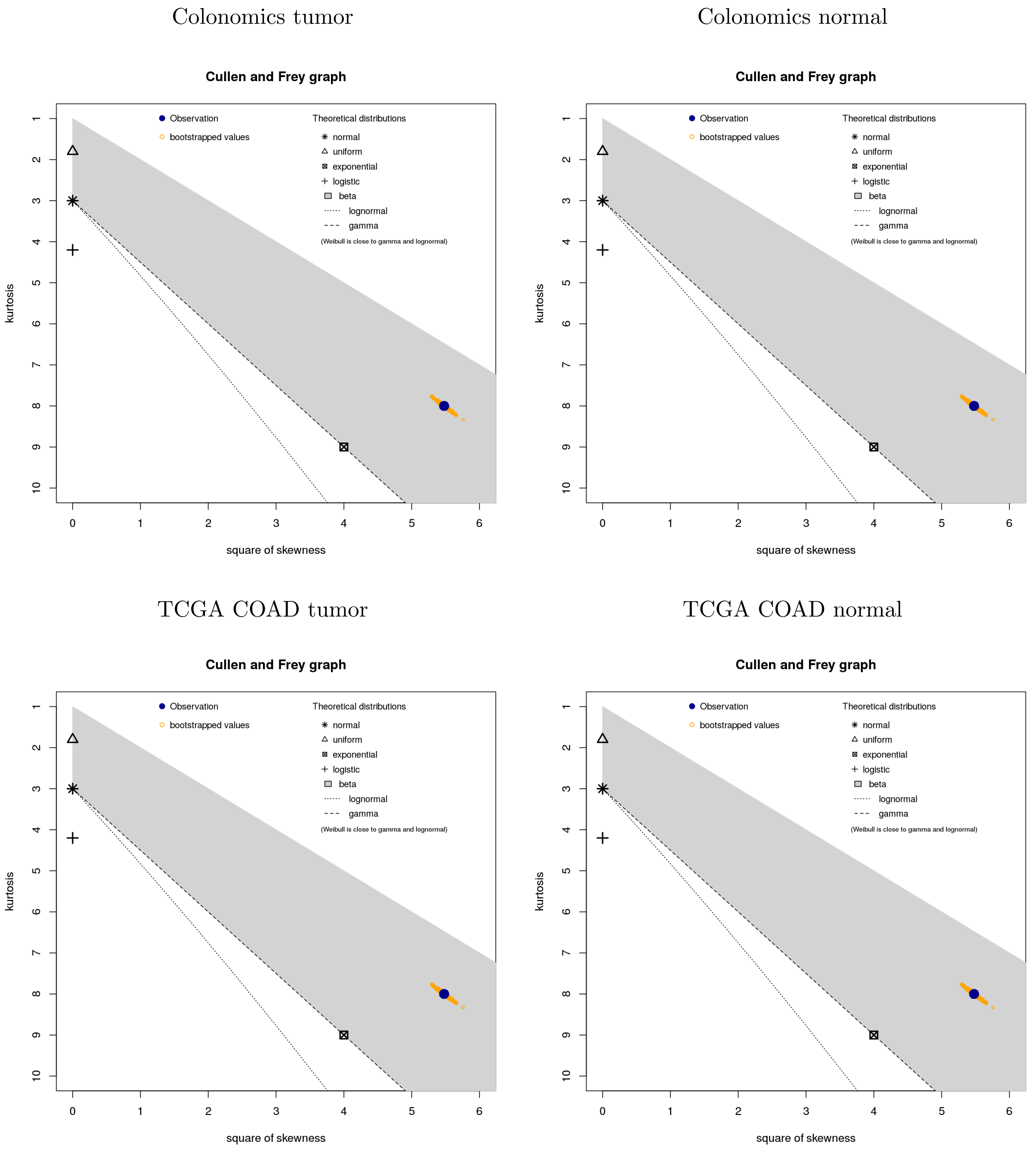

Figure S8: Cullen and Frey graphs. Cullen and Frey graph depicting the skewness-kurtosis plot of the network degree distribution (in blue) for each dataset, pointing to differences to the expected values for some theoretical distributions (i.e. lognormal, exponential, gamma) (Cullen and Frey, 1999; Delignette-Muller and Dutang, 2015).

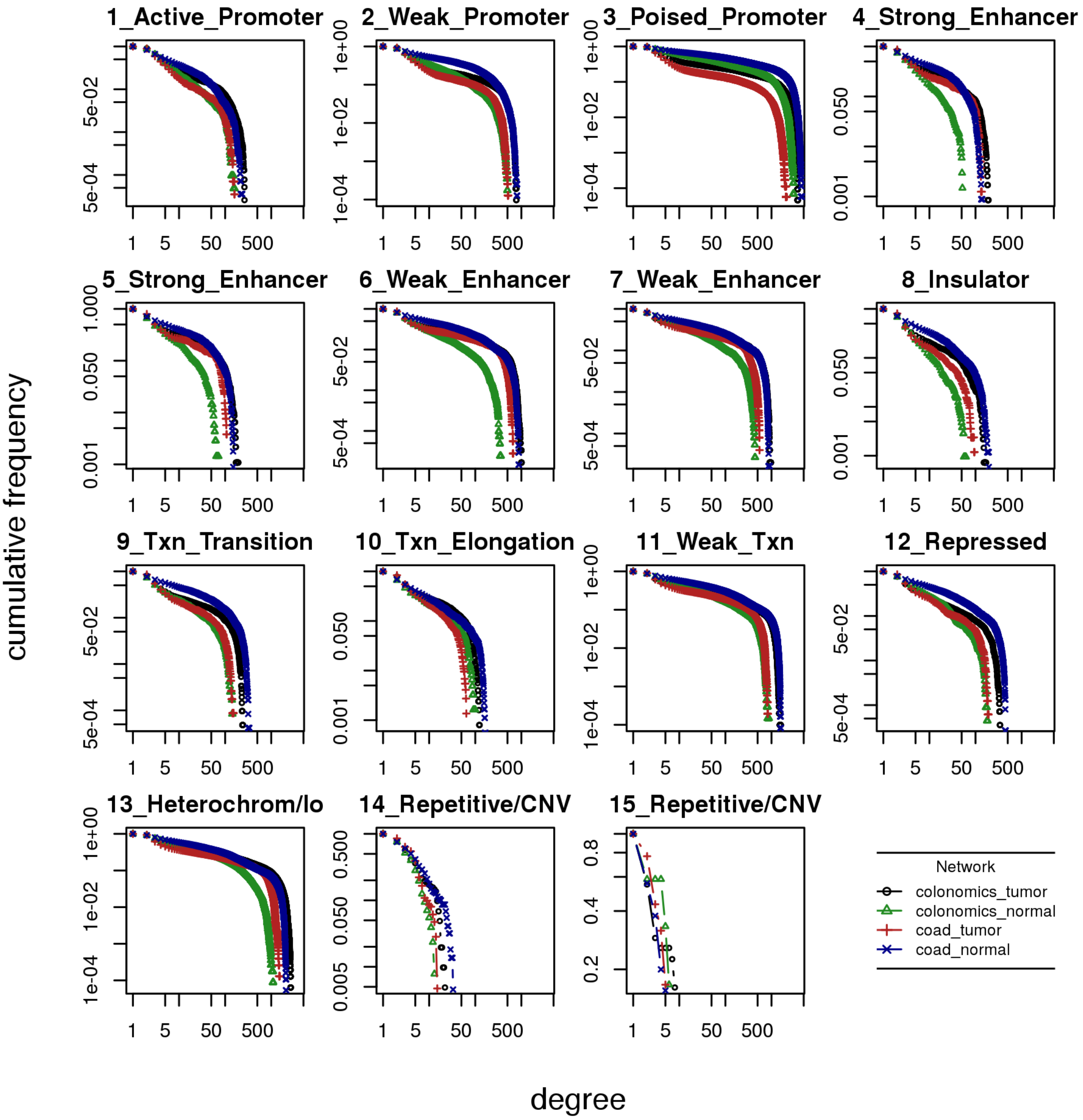

Figure S9: Complementary cumulative distribution function of node degrees stratified by chromatin color (15-states HMM as predicted for human H1 stem cells).

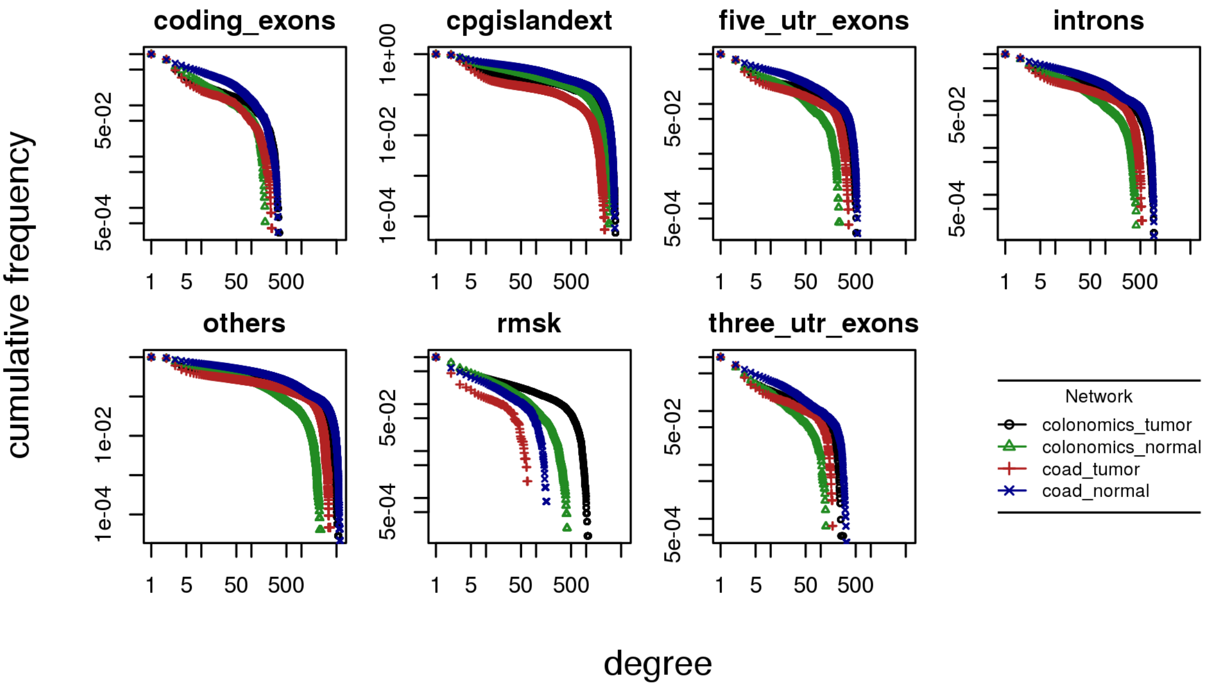

Figure S10: Complementary cumulative distribution function of node degrees stratified by genomic compartment. cpgislandext: CpG island; rmsk: repeats as predicted by RepeatMasker; coding exons, three utr and five utr as described by the Ensembl ensGene table.

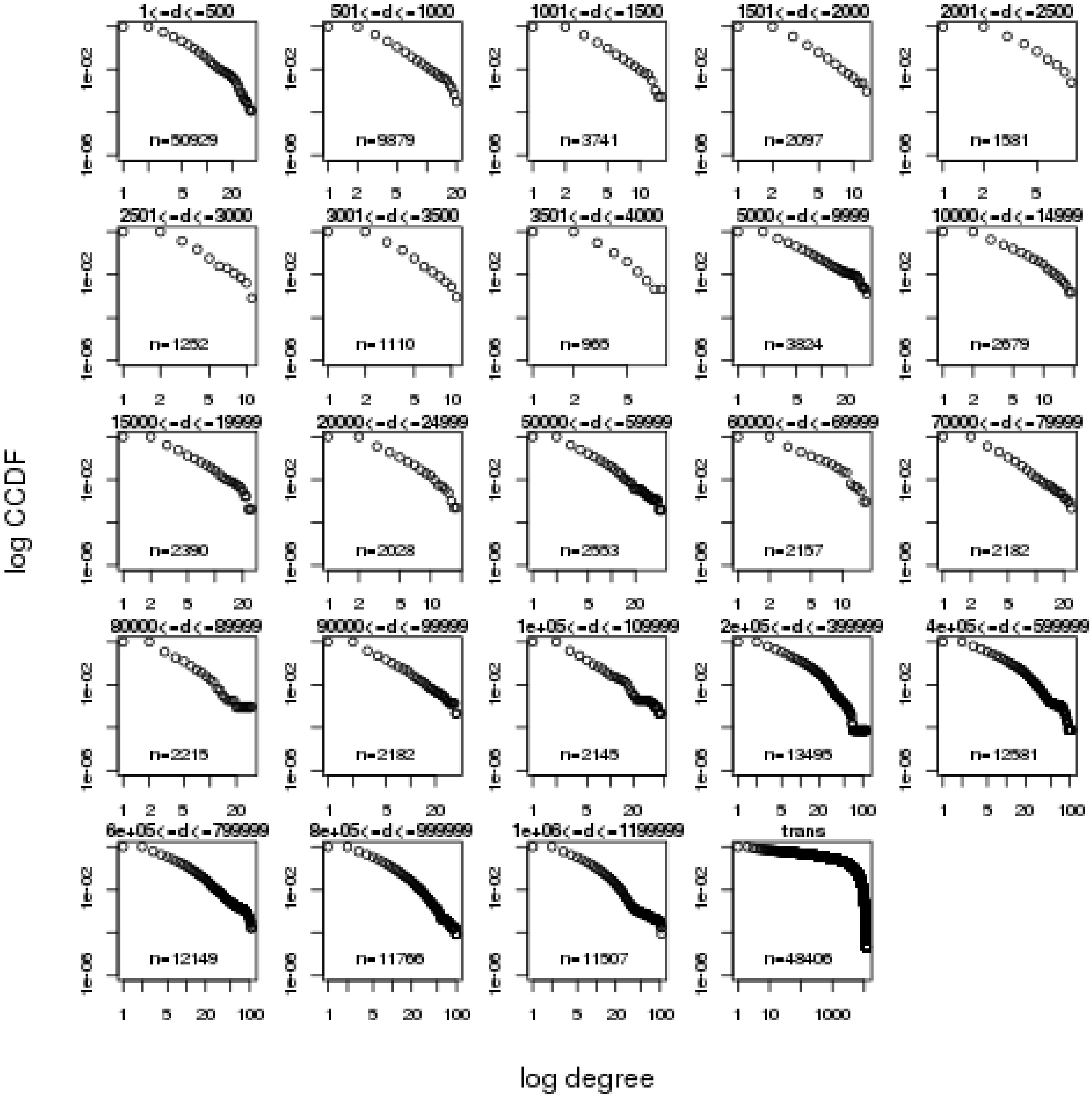

Figure S11: Complementary cumulative distribution function (ccdf) vs node degree (log-log transformed) for the colonomics tumor cohort. Stratified by distance between probes.

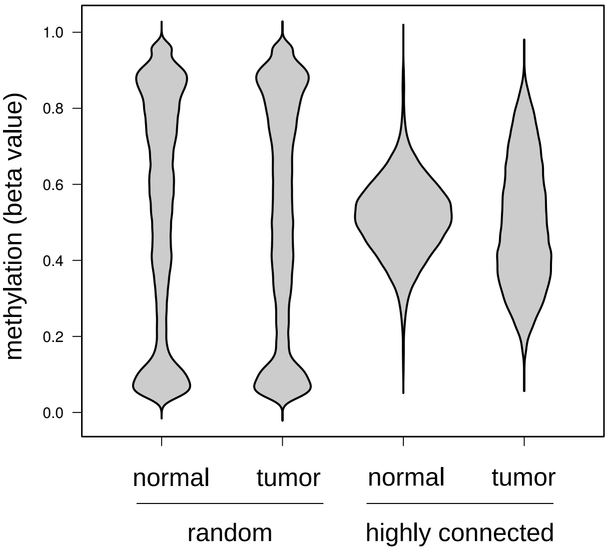

Figure S12: Top connectivity probes DNA methylation distribution in Colonomics tumor and Colonomics normal as compared to a subset of random CpGs. Sorting by connectivity was defined according to each probe's degree (number of different co-methylation partners); we consider the 99% percentile (that is, the 1% richest probes) as the top connected ones. An equally sized background was retrieved by random sampling the Infinium 450k array. DNA methylation (beta value) data was retrieved from both Colonomics tumor and Colonomics normal for each set of probes, the top connected in tumor and the random background.

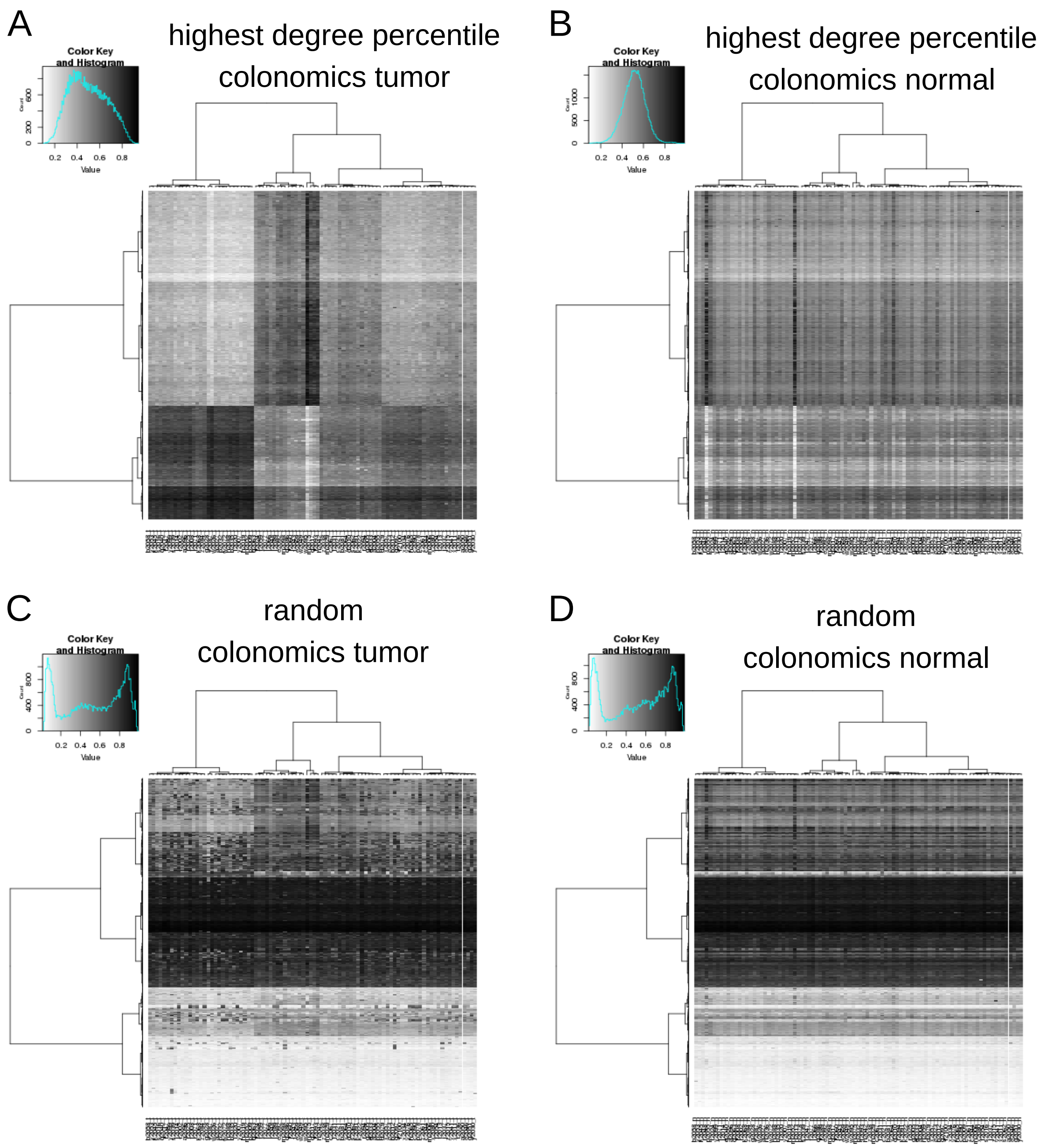

Figure S13: Top connectivity probes DNA methylation profiles (as defined in figure S12) in Colonomics tumor (A) and Colonomics normal (B). As compared to a subset of random CpGs (C and D), Colonomics tumor show heterogeneous DNA methylation structure (as expected, driving co-methylations) whereas normals show homogeneous intermediate values. Samples (columns) and CpGs (rows) are sorted as in (A).

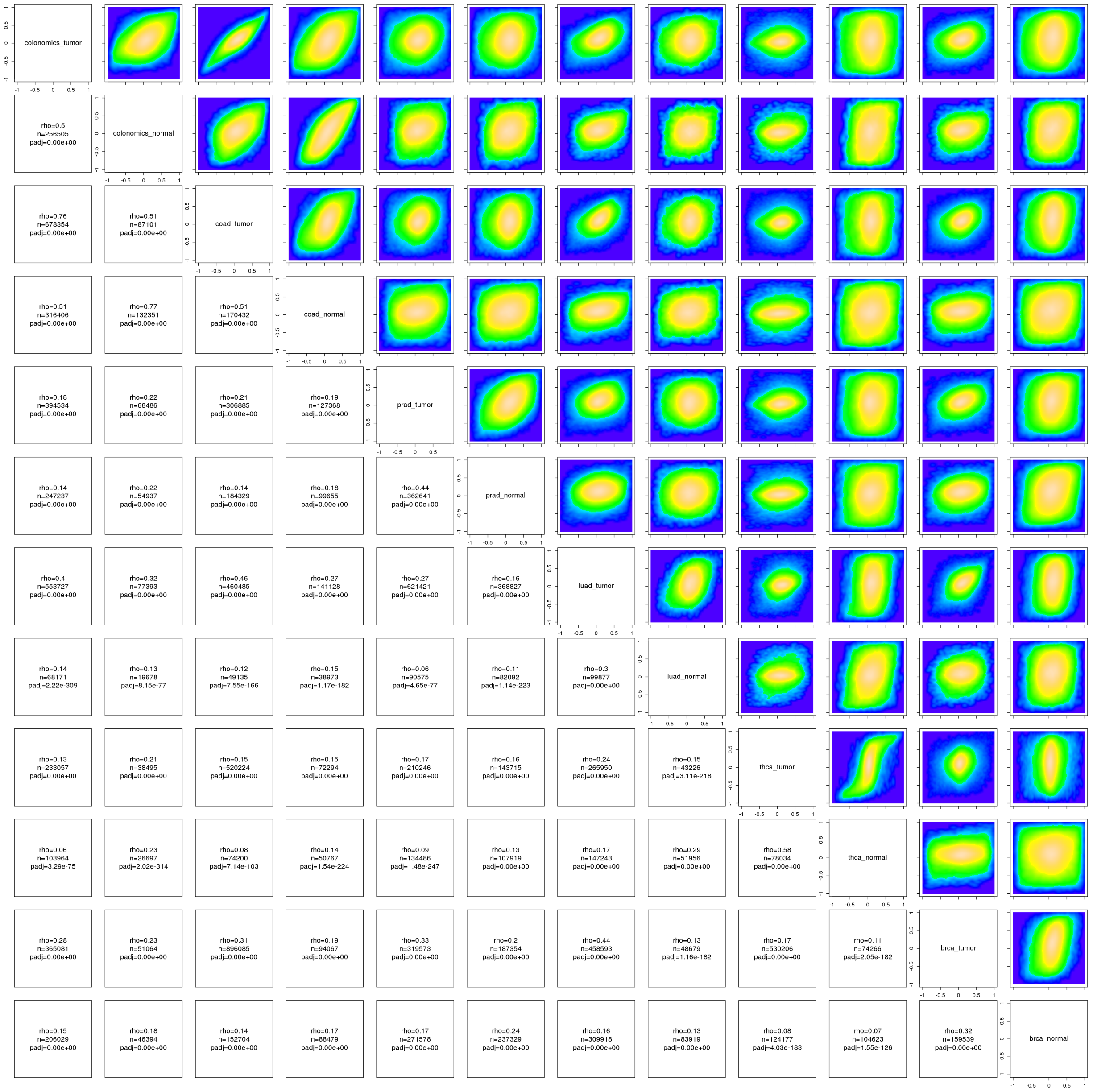

Figure S14: Comethylation landscape is tumor type-specific. Comparison of the DNA co-methylation distribution across TCGA types only depicts sharedness between TCGA colon and the original colon datasets, indicating the co-methylation landscape is tumor-type specific.

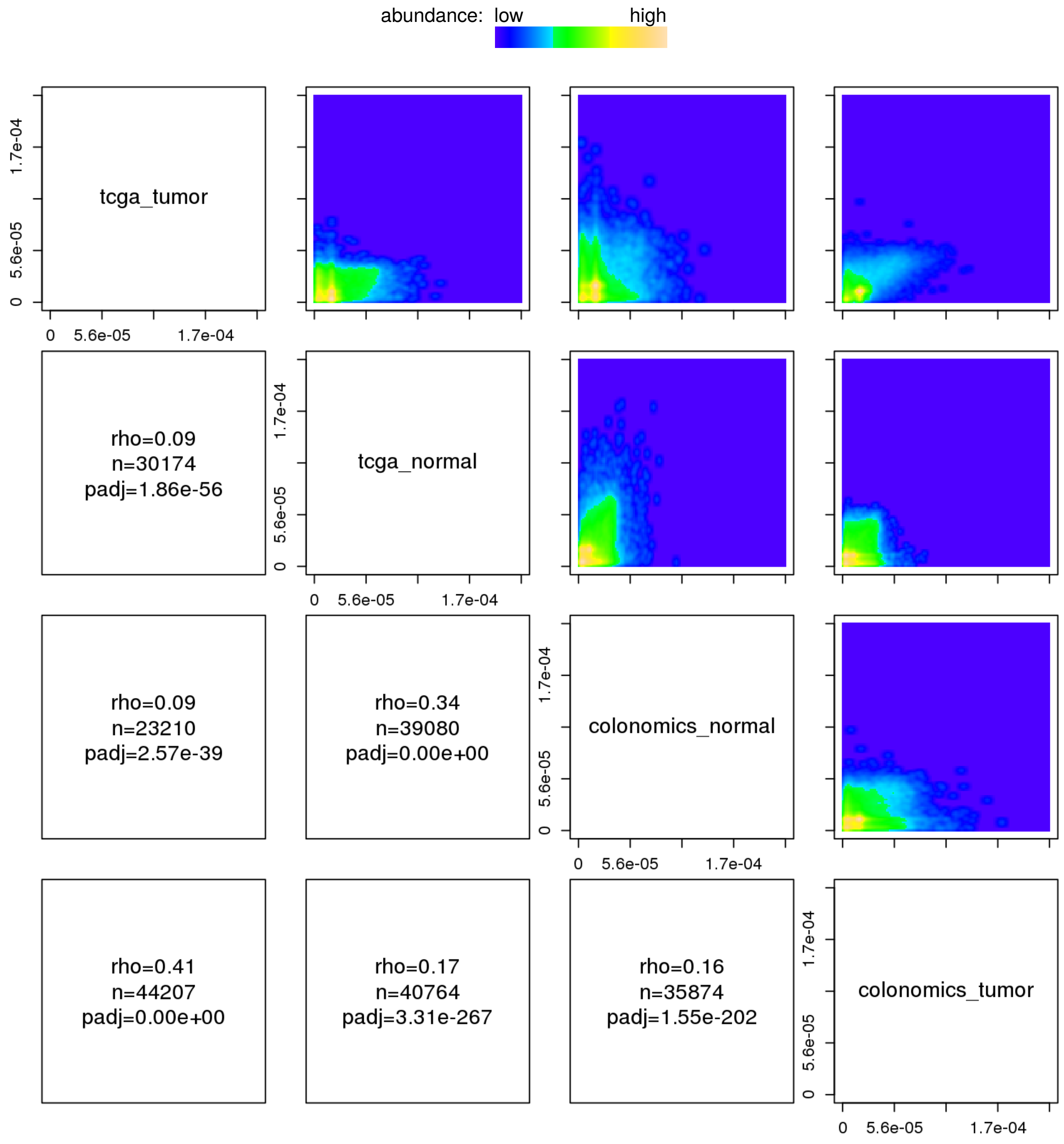

Figure S15: Pagerank score (influence) correspondence across cohorts. Pairwise comparisons face nodes (CpGs) present at both cohorts, as sampled from each individual network ($\rho >= 0.8$). X and Y axis depict Pagerank values.

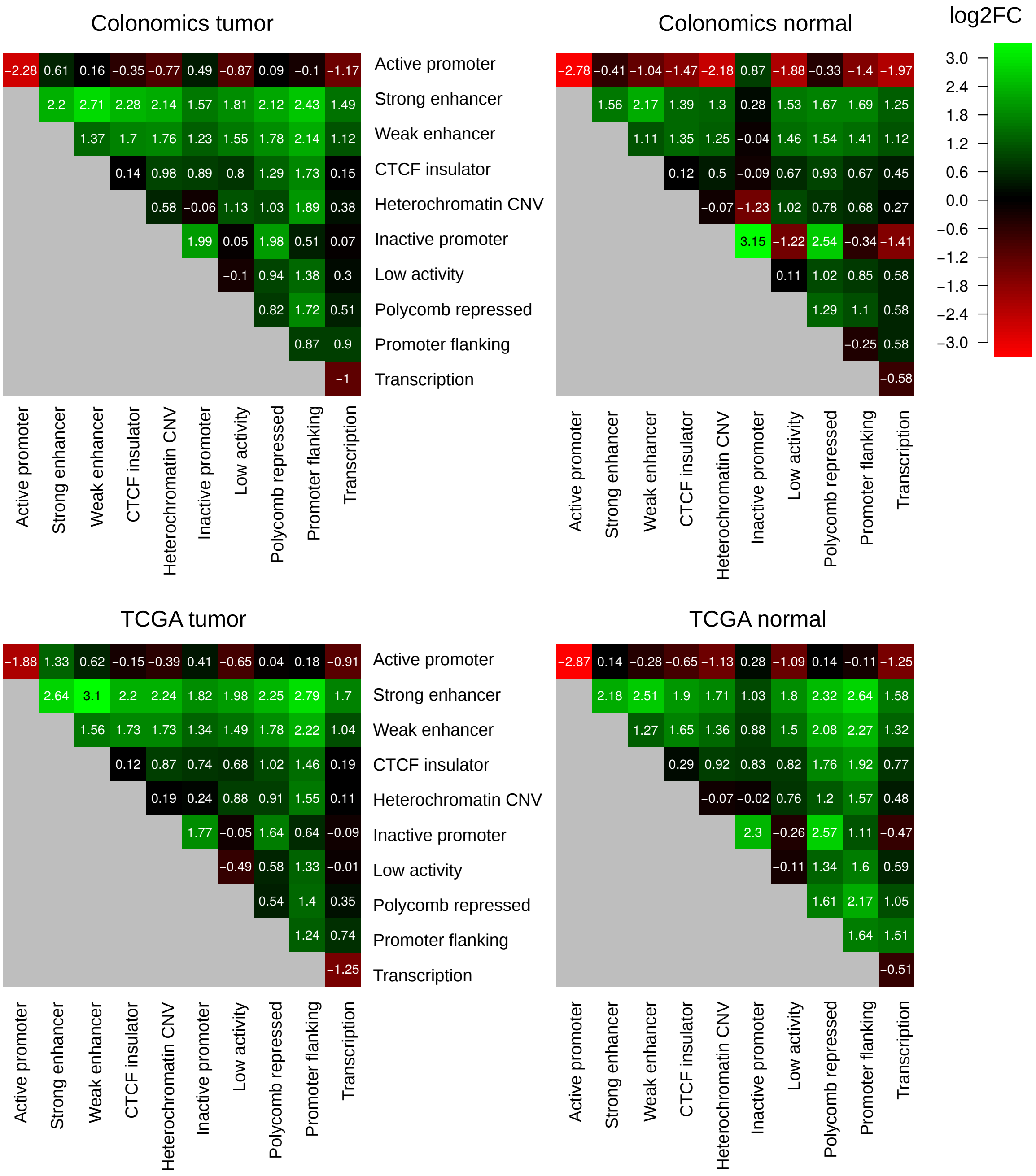

Figure S16: Pairwise chromatin color co-methylation enrichment (log2 fold change as compared to the Infinium chip background).

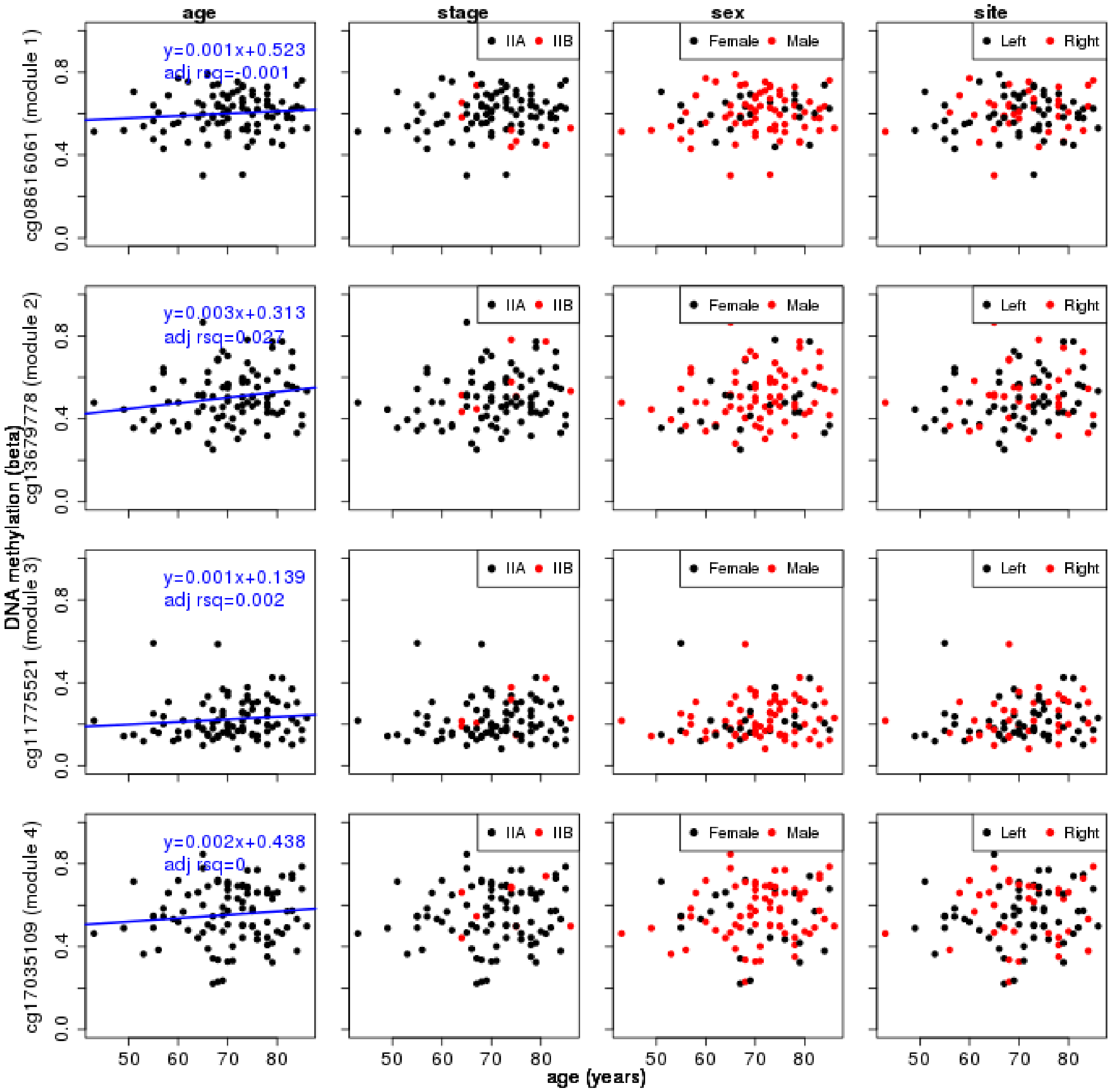

Figure S17: Modules methylation is not driven by age. The DNA methylation status of a randomly picked CpG representative from each of the top 4 modules are not strongly associated to age, disease stage, gender nor anatomical site. Dots represent primary tumors from the Colonomics cohort sorted by the patient's age.

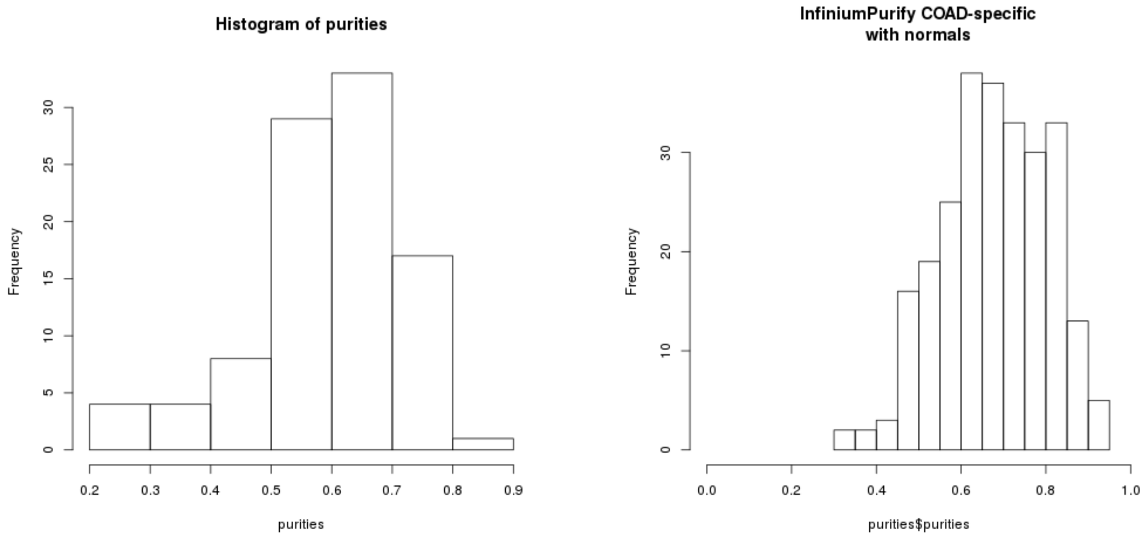

Figure S18: Purity histograms as called by the InfiniumPurify method at the Colonomics (left) and TCGA (right) datasets.

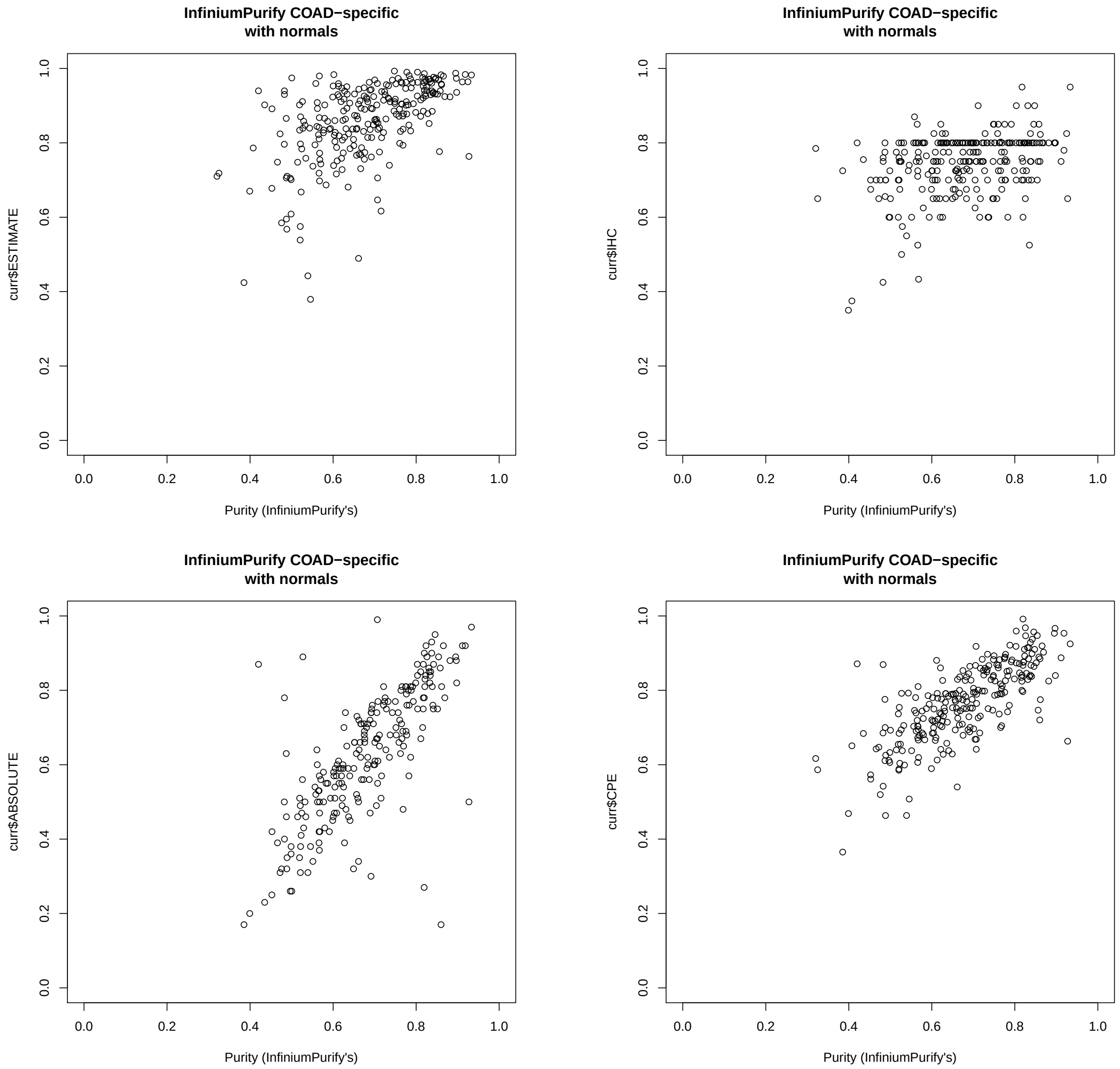

Figure S19: InfiniumPurify purity estimates based on DNA methylation data. A local run of InfiniumPurify on TCGA colon tumors is compared to the ESTIMATE, ABSOLUTE, IHC and CPE indices as calculated by the TCGA meta-analysis by Aran et al. (2015).

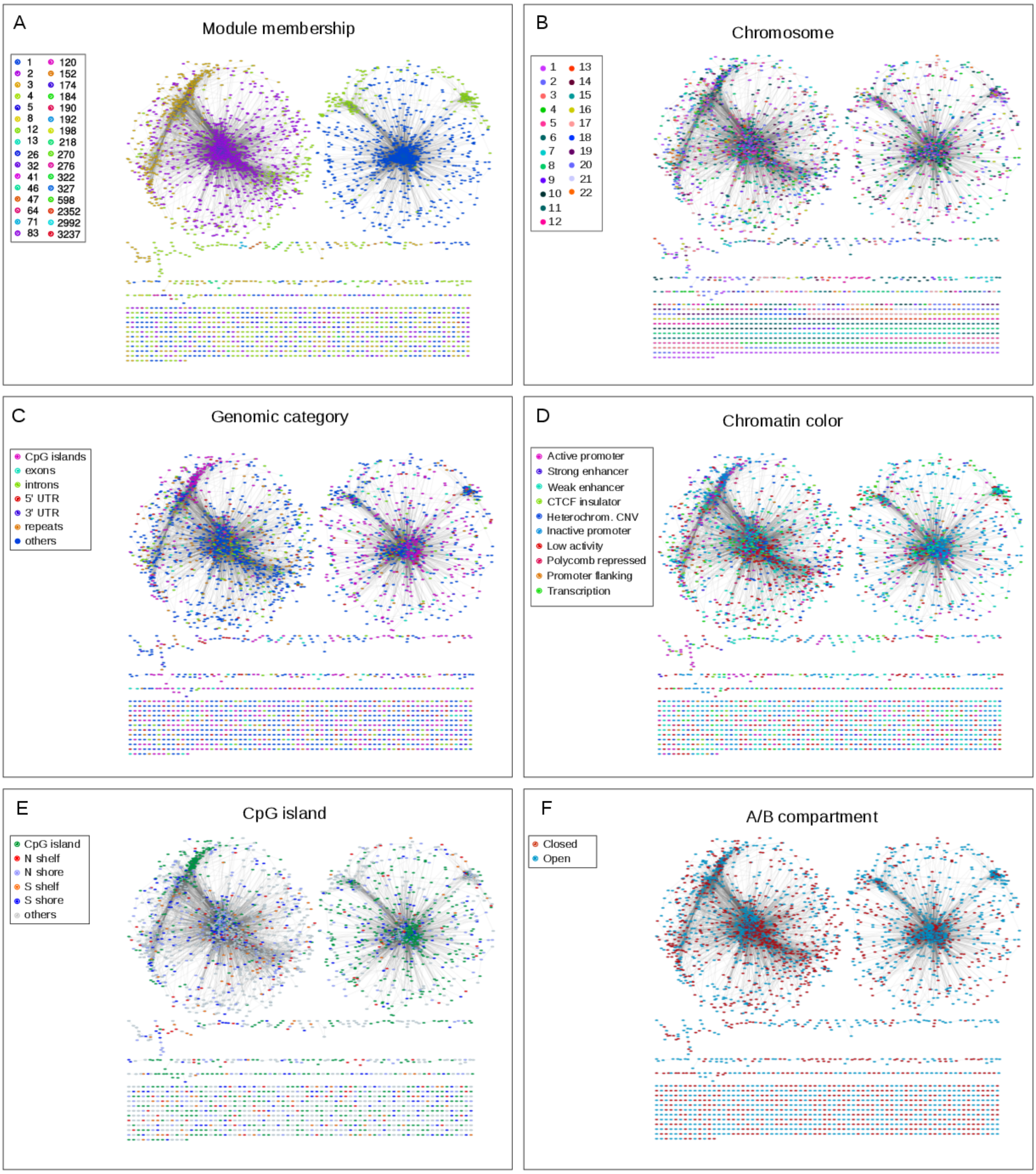

Figure S20: Co-methylation network modularity. **A**, The tumor co-methylation network (Colonomics tumor) is modular and contains giant components. Genomic loci (network nodes) coloring: **B**, chromosome; **C**, genomic category; **D**, chomatin color; **E**, location to CpG islands; and **F**, open/closed compartment. Graphs depicted here are limited to a random sample of 5000 nodes located at trans modules; network layout was calculated by $1 - \rho$ (edges) weighted springs.

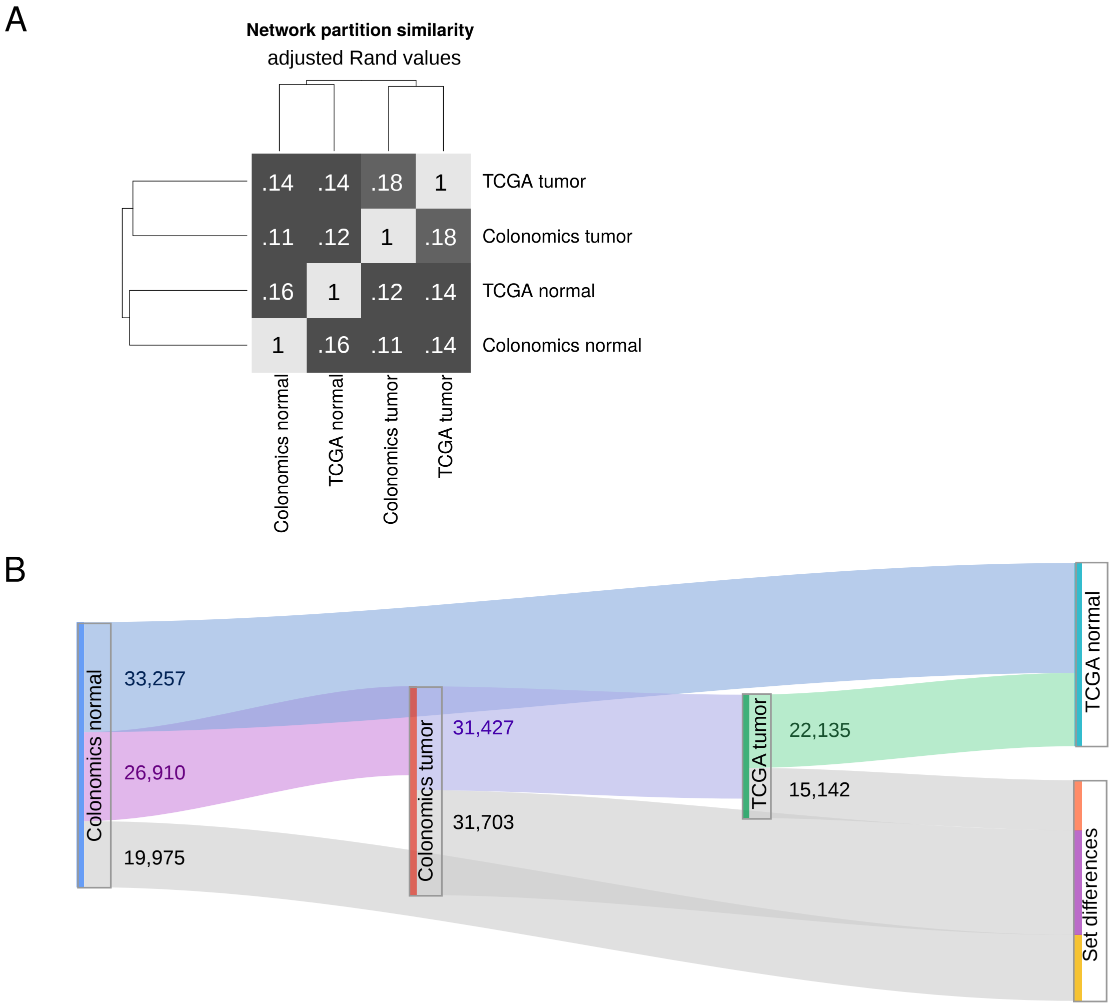

Figure S21: Datasets comparison (Adjusted Rand Indices). **A**, adjusted Rand similarities built upon significant CpGs ($\rho \geq 0.8$). **B**, Sankey diagram depicting the number of shared CpGs across datasets.

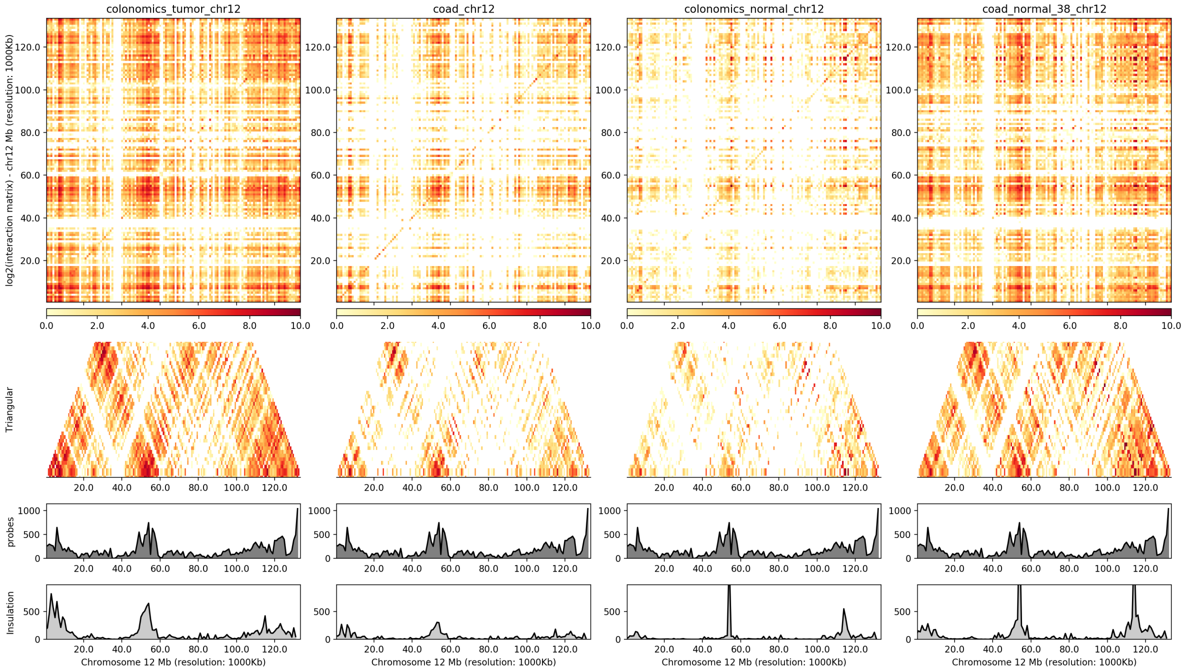

Figure S22: HiC-like cross cohort visualization. The number of correlations between 1 Mbp-long bins of chromosome 12 were represented as a HiC contact matrix. Tracks depict a triangular view of the whole matrix, the Infinium 450k probe density and the insulation score from the cis-comethylating probes. From left to right: Colonomics tumor, TCGa tumor, Colonomics normal, TCGA normal.

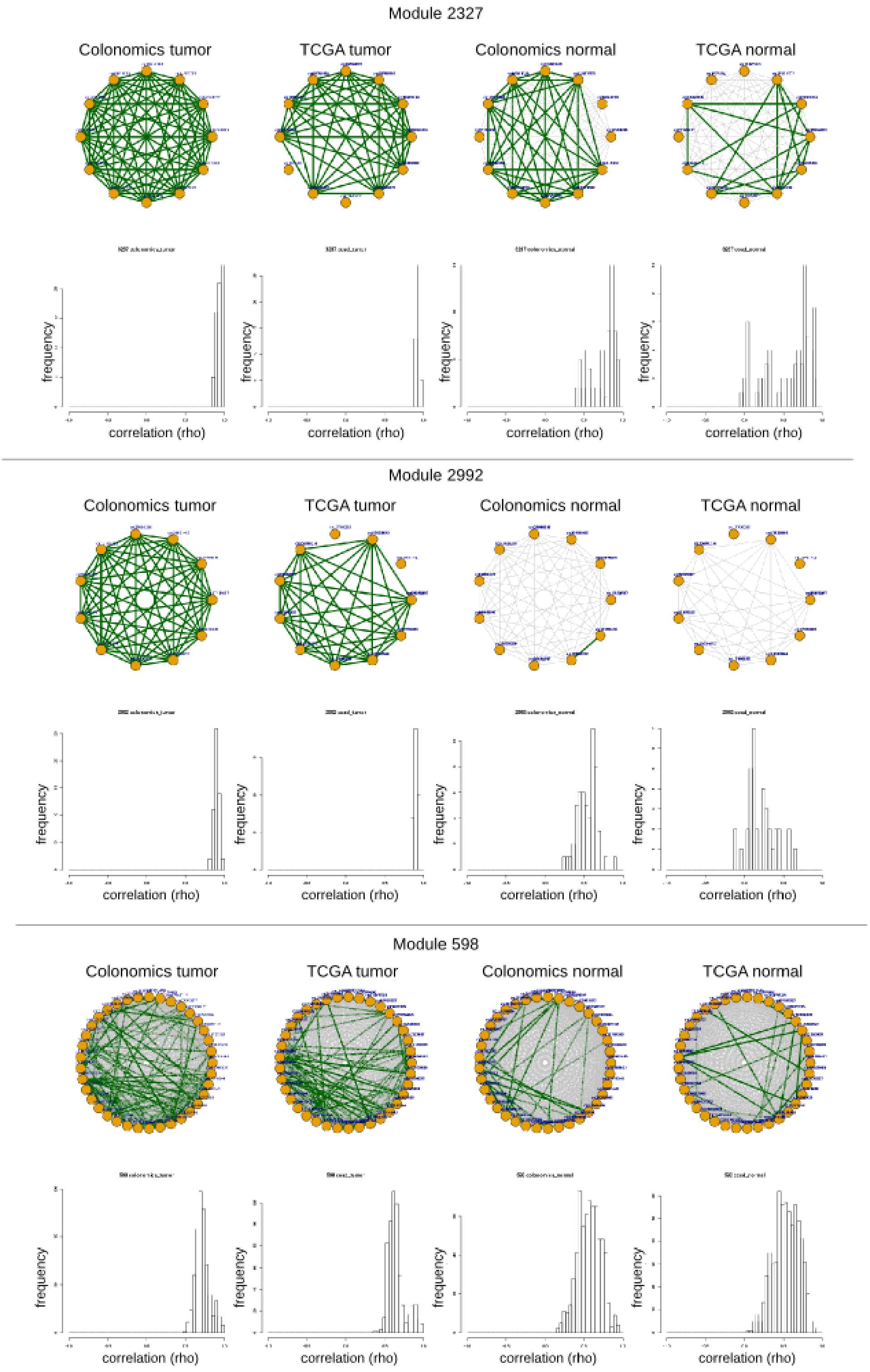

Figure S23: Co-methylation modules conservation. Significant correlations ($\rho \geq 0.8$) are depicted as green links. Some CpGs are not available for some cohorts due to quality filters (see supplementary methods).

| | colonomics_tumor 1 (15259) | colonomics_tumor 2 (18727) | colonomics_tumor 3 (5099) | colonomics_tumor 4 (6750) | colonomics_tumor 5 (385) | colonomics_tumor 8 (168) | colonomics_tumor 12 (31) | colonomics_tumor 174 (16) | colonomics_tumor 192 (36) | colonomics_tumor 322 (16) | colonomics_tumor 598 (37) | colonomics_tumor 2352 (32) |
|---|---|---|---|---|---|---|---|---|---|---|---|---|
| *colonomics_normal 1 (17758)* | 1971<br>1 | 1763<br>1 | 567<br>3.83e−05 | 1506<br>0 | 0<br>1 | 0<br>1 | 0<br>1 | 0<br>1 | 0<br>1 | 0<br>1 | 0<br>1 | 0<br>1 |
| *colonomics_normal 2 (9299)* | 5274<br>0 | 0<br>1 | 1<br>1 | 2<br>1 | 56<br>5.91e−06 | 0<br>1 | 0<br>1 | 10<br>2.77e−06 | 0<br>1 | 0<br>1 | 2<br>0.978 | 5<br>0.41 |
| *colonomics_normal 3 (9342)* | 26<br>1 | 5434<br>0 | 358<br>1 | 190<br>1 | 0<br>1 | 0<br>1 | 0<br>1 | 0<br>1 | 0<br>1 | 0<br>1 | 0<br>1 | 0<br>1 |
| *colonomics_normal 4 (3404)* | 180<br>1 | 588<br>1 | 713<br>0 | 12<br>1 | 63<br>1.68e−36 | 0<br>1 | 0<br>1 | 0<br>1 | 15<br>6.53e−17 | 12<br>1.14e−13 | 16<br>7.05e−16 | 10<br>3.34e−08 |
| *colonomics_normal 5 (369)* | 101<br>3.05e−29 | 6<br>1 | 1<br>1 | 3<br>0.998 | 0<br>1 | 0<br>1 | 0<br>1 | 0<br>1 | 0<br>1 | 0<br>1 | 0<br>1 | 0<br>1 |
| *colonomics_normal 8 (472)* | 0<br>1 | 13<br>1 | 0<br>1 | 8<br>0.456 | 0<br>1 | 60<br>1.16e−156 | 0<br>1 | 0<br>1 | 0<br>1 | 0<br>1 | 0<br>1 | 0<br>1 |
| *colonomics_normal 14 (20)* | 14<br>2.95e−06 | 0<br>1 | 0<br>1 | 0<br>1 | 0<br>1 | 0<br>1 | 0<br>1 | 0<br>1 | 0<br>1 | 0<br>1 | 0<br>1 | 0<br>1 |
| *colonomics_normal 24 (22)* | 20<br>1.26e−08 | 0<br>1 | 0<br>1 | 0<br>1 | 0<br>1 | 0<br>1 | 0<br>1 | 0<br>1 | 0<br>1 | 0<br>1 | 0<br>1 | 0<br>1 |
| *colonomics_normal 39 (17)* | 14<br>2.95e−06 | 0<br>1 | 0<br>1 | 0<br>1 | 0<br>1 | 0<br>1 | 0<br>1 | 0<br>1 | 0<br>1 | 0<br>1 | 0<br>1 | 0<br>1 |
| *colonomics_normal 97 (66)* | 0<br>1 | 0<br>1 | 0<br>1 | 0<br>1 | 0<br>1 | 0<br>1 | 29<br>5.31e−94 | 0<br>1 | 0<br>1 | 0<br>1 | 0<br>1 | 0<br>1 |
| *colonomics_normal 664 (21)* | 18<br>5.65e−06 | 0<br>1 | 0<br>1 | 2<br>0.553 | 0<br>1 | 0<br>1 | 0<br>1 | 0<br>1 | 0<br>1 | 0<br>1 | 0<br>1 | 0<br>1 |
| *colonomics_normal 679 (12)* | 12<br>1.82e−05 | 0<br>1 | 0<br>1 | 0<br>1 | 0<br>1 | 0<br>1 | 0<br>1 | 0<br>1 | 0<br>1 | 0<br>1 | 0<br>1 | 0<br>1 |
| *colonomics_normal 697 (10)* | 0<br>1 | 10<br>0.000123 | 0<br>1 | 0<br>1 | 0<br>1 | 0<br>1 | 0<br>1 | 0<br>1 | 0<br>1 | 0<br>1 | 0<br>1 | 0<br>1 |
| *colonomics_normal 896 (12)* | 1<br>0.994 | 0<br>1 | 0<br>1 | 9<br>3.77e−09 | 0<br>1 | 0<br>1 | 0<br>1 | 0<br>1 | 0<br>1 | 0<br>1 | 0<br>1 | 0<br>1 |
| *colonomics_normal 925 (16)* | 12<br>1.82e−05 | 0<br>1 | 0<br>1 | 0<br>1 | 0<br>1 | 0<br>1 | 0<br>1 | 0<br>1 | 0<br>1 | 0<br>1 | 0<br>1 | 0<br>1 |
| *colonomics_normal 1033 (13)* | 13<br>7.34e−06 | 0<br>1 | 0<br>1 | 0<br>1 | 0<br>1 | 0<br>1 | 0<br>1 | 0<br>1 | 0<br>1 | 0<br>1 | 0<br>1 | 0<br>1 |
| *colonomics_normal 1212 (32)* | 31<br>5.59e−13 | 0<br>1 | 0<br>1 | 0<br>1 | 0<br>1 | 0<br>1 | 0<br>1 | 0<br>1 | 0<br>1 | 0<br>1 | 0<br>1 | 0<br>1 |
| *colonomics_normal 1217 (14)* | 12<br>1.82e−05 | 0<br>1 | 0<br>1 | 0<br>1 | 0<br>1 | 0<br>1 | 0<br>1 | 0<br>1 | 0<br>1 | 0<br>1 | 0<br>1 | 0<br>1 |
| *colonomics_normal 1221 (15)* | 10<br>0.000112 | 0<br>1 | 0<br>1 | 0<br>1 | 0<br>1 | 0<br>1 | 0<br>1 | 0<br>1 | 0<br>1 | 0<br>1 | 0<br>1 | 0<br>1 |
| *colonomics_normal 1224 (13)* | 11<br>4.53e−05 | 0<br>1 | 0<br>1 | 0<br>1 | 0<br>1 | 0<br>1 | 0<br>1 | 0<br>1 | 0<br>1 | 0<br>1 | 0<br>1 | 0<br>1 |
| *colonomics_normal 1241 (43)* | 26<br>0.000778 | 0<br>1 | 0<br>1 | 13<br>2.31e−05 | 0<br>1 | 0<br>1 | 0<br>1 | 0<br>1 | 0<br>1 | 0<br>1 | 0<br>1 | 0<br>1 |

Figure S24: Network partitioning overlap between colon tumor and normal. Module preservation across cohorts was calculated by cross-tabulating the CpGs belonging to each community (Langfelder et al., 2011).

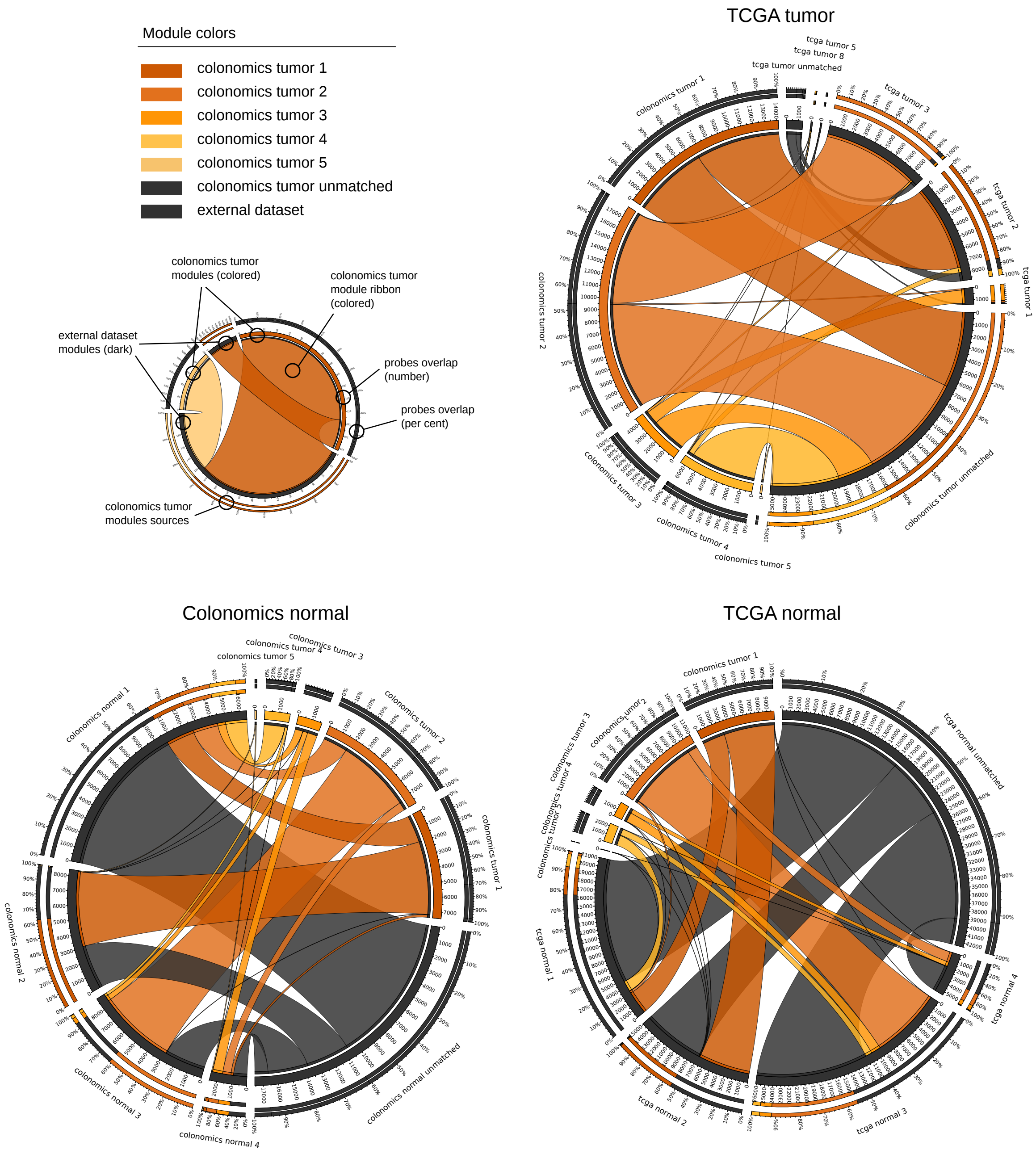

Figure S25: Equivalences to the top 5 Colonomics tumor communities. The CpGs from the top 5 Colonomics tumor communities partly overlap to TCGA tumor, Colonomics normal and TCGA normal communities (see Figure S24 for a comprehensive overview). Concentric tracks depict the number of CpGs involved (relative and absolute values for outer and inner tracks, respectively). Ribbons are colored according to the Colonomics tumor community. Segments tagged as 'unmatched' depict CpGs that correlate with $\rho \geq 0.8$ at a single dataset.

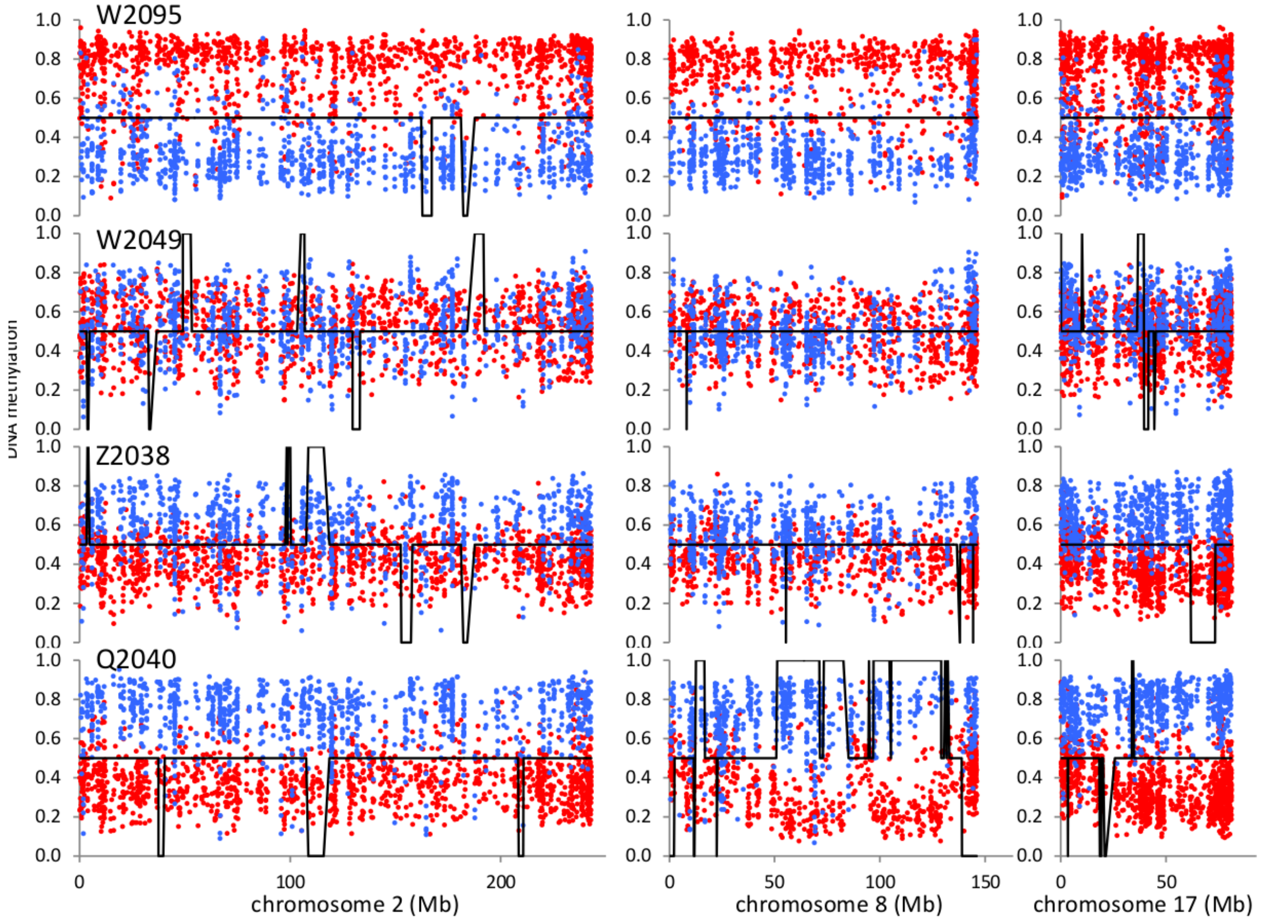

Figure S26: DNA methylation and copy number profiles in chromosomes 2, 8 and 17 in four tumor samples (W2095, W2049, Z2038 and Q2040). DNA methylation beta values of CpGs belonging to modules 1 (blue) and 2 (red) are displayed as dots. Chromosomal imbalances analyzed by array CGH are shown as low (losses) or high (gains) segments along the centered black line representing balanced chromosome segments.

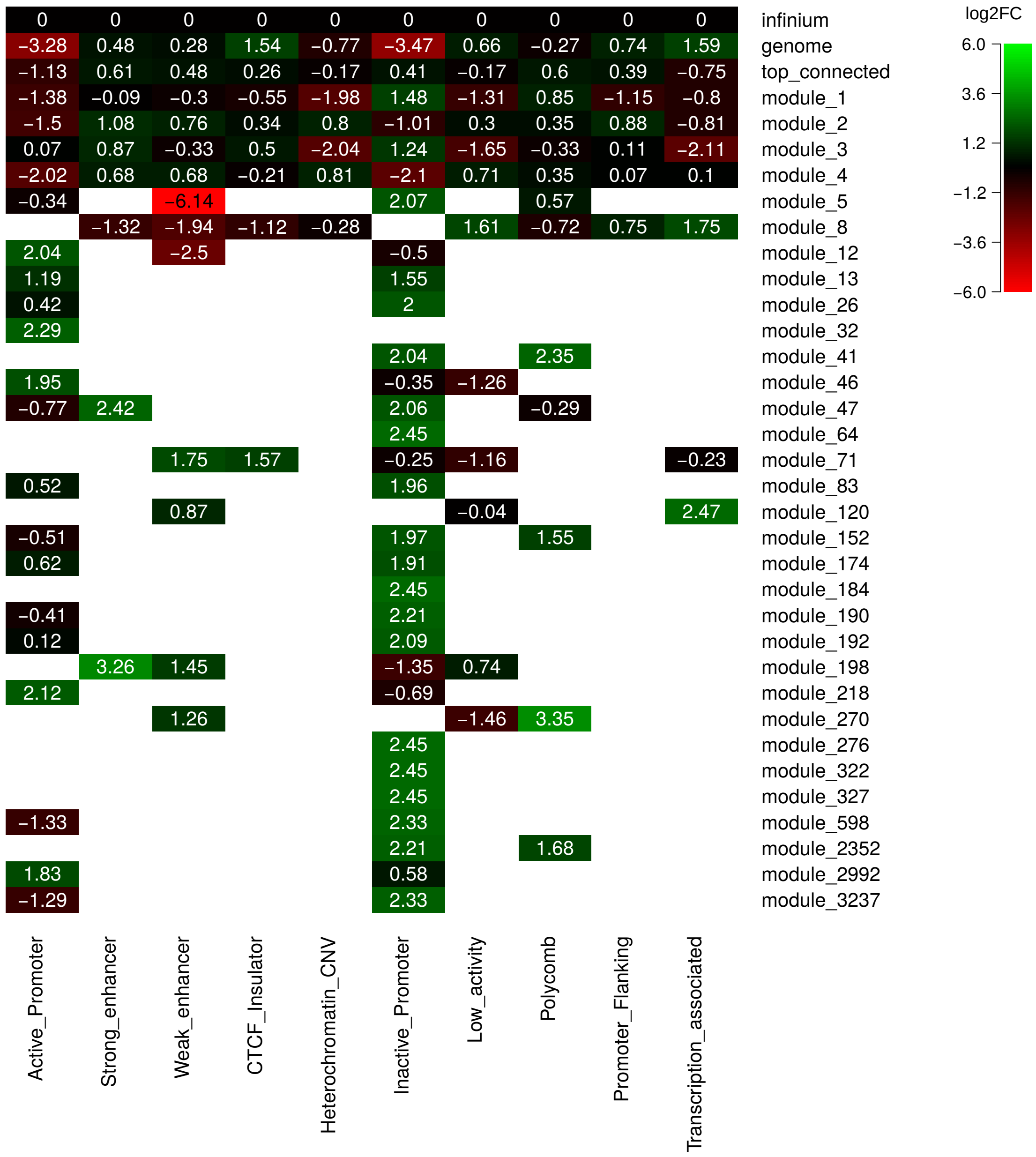

Figure S27: HMM colors enrichment for the top connected probes (the ones at the 99th percentile of node degree) and each module's. For each row, the proportion of probes annotated to each HMM state (column) is normalized to the Infinium background. Colors depict log2 fold changes (log2FC) depict enrichment (positive, green), no change (zero, black) or depletion (negative, red) as compared to the Infinium chip. Importantly, the Infinium background is not a fair representation of the genome (first row).

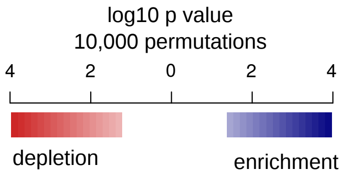

| hypo | hyper | boundary shift | loss of regulation | novel methylation | other | |
|---|---|---|---|---|---|---|
| 4 | 4 | 4 | 4 | 0.47 | 4 | module_1 |
| 4 | 1.15 | 4 | 0.87 | 4 | 4 | module_2 |
| 4 | 0.28 | 4 | 1.07 | 1.97 | 4 | module_3 |
| 4 | 4 | 4 | 4 | 4 | 4 | module_4 |
| 4 | 1.87 | 4 | 4 | 0.01 | 4 | module_5 |
| 4 | 1.27 | 0.05 | 0.92 | 0.01 | 0.1 | module_8 |
| 4 | 0.18 | 0.01 | 0.01 | 0 | 0.02 | module_12 |
| 0.77 | 0.01 | 3.52 | 3.1 | 0 | 0 | module_13 |
| 1.2 | 0.02 | 3.1 | 0 | 0 | 4 | module_26 |
| 0.46 | 0.01 | 4 | 0 | 0 | 0 | module_32 |
| 0.73 | 0.02 | 0 | 0 | 0 | 4 | module_41 |
| 1.89 | 1.55 | 0 | 2.68 | 0 | 2.21 | module_46 |
| 3.3 | 0.05 | 2.92 | 2.68 | 0 | 4 | module_47 |
| 1.12 | 1.59 | 0 | 0 | 0 | 2.96 | module_64 |
| 1.81 | 0.06 | 0 | 0 | 0 | 1.76 | module_71 |
| 0.85 | 1.56 | 3.52 | 0 | 0 | 3.7 | module_83 |
| 4 | 0.12 | 0.01 | 0.01 | 0 | 0.01 | module_120 |
| 2.17 | 0.02 | 2.92 | 2.62 | 0 | 3.3 | module_152 |
| 1.47 | 0.01 | 0 | 2.47 | 0 | 4 | module_174 |
| 1.39 | 0.02 | 3.1 | 0 | 0 | 2.49 | module_184 |
| 1.32 | 1.74 | 3.7 | 4 | 0 | 2.48 | module_190 |
| 2.4 | 0.05 | 3.7 | 0 | 0 | 0.01 | module_192 |
| 0.41 | 0.03 | 2.89 | 0 | 0 | 2.31 | module_198 |
| 1.33 | 0.04 | 0 | 0 | 0 | 0.01 | module_218 |
| 1.03 | 0.01 | 0 | 0 | 0 | 0 | module_270 |
| 1.04 | 0.01 | 0 | 0 | 0 | 3.15 | module_276 |
| 1.07 | 0.02 | 0 | 4 | 0 | 0 | module_322 |
| 1.03 | 0.01 | 2.68 | 0 | 0 | 0 | module_327 |
| 3 | 0.05 | 0 | 4 | 0 | 1.93 | module_598 |
| 1.32 | 0.01 | 0 | 4 | 0 | 2.41 | module_2352 |
| 0.77 | 0.02 | 0 | 2.96 | 0 | 0 | module_2992 |
| 0.98 | 0.01 | 3.15 | 0 | 0 | 0 | module_3237 |

Figure S28: Statistical association to known variably methylated regions (Hansen et al., 2011). We evaluated positive association (in blue) and anti-association (in red) between them using a permutation approach (Gel et al., 2015) using the Infinium 450k CpGs with enough variability in our cohort as background. As described in (Hansen et al., 2011), DMRs include tumor hypo- and hypermethylations; CpG island methylation boundaries, e.g. boundary shifts (spreading of CpG island methylation into adjacent regions) and loss of regulation (loss of boundary sharpness); and novel hypomethylation events. Color scale indicates log10-transformed pvalues after 10,000 permutations (white: no significance, Z-scores at table S7).

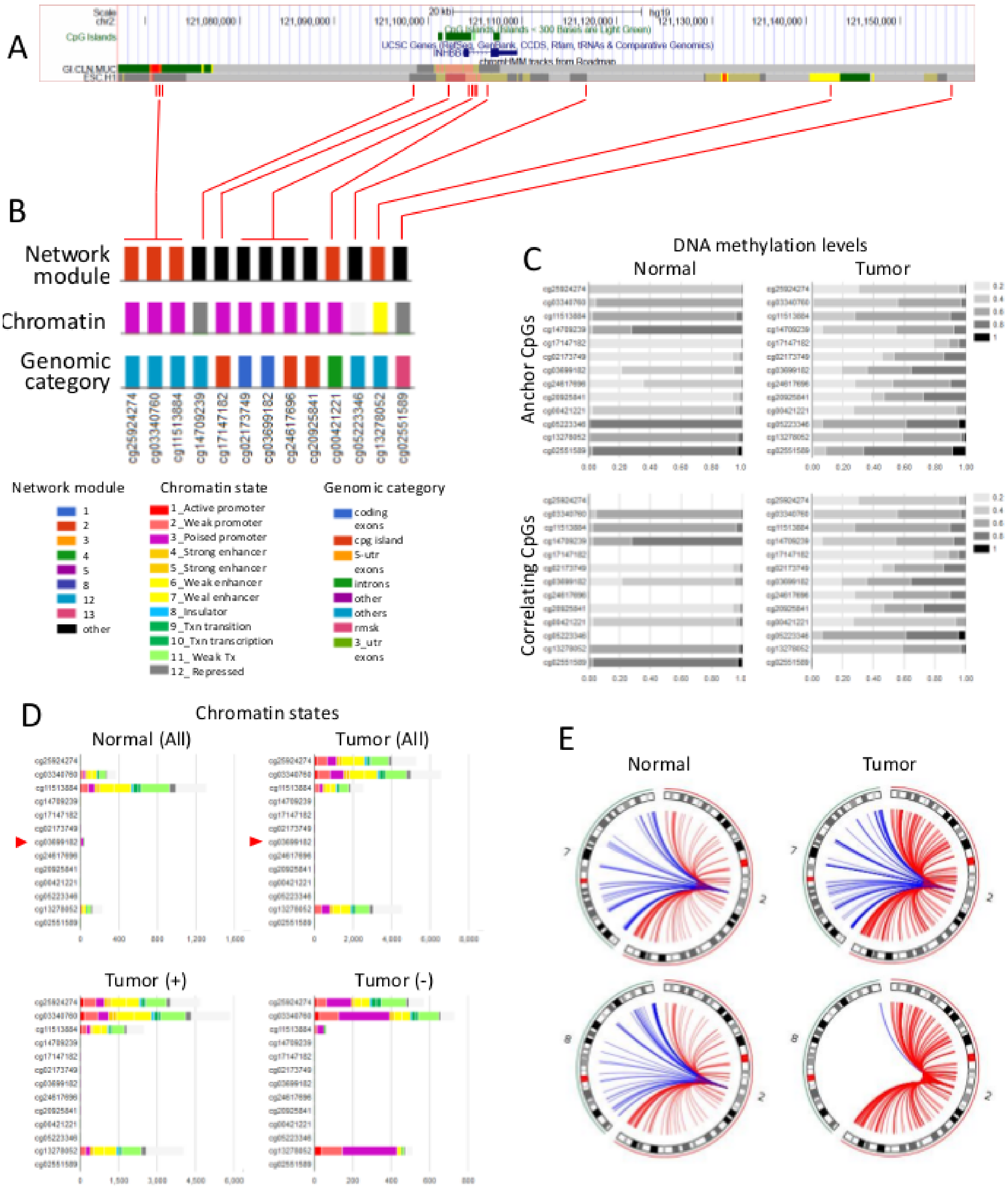

Figure S29: *Corre* web tool usage when selecting a CpG (i.e.: cg25924274) or a set of gene associated probes (i.e.: INHBB). Corre renders graphs displaying diverse features of the co-methylating sites, including DNA methylation levels, genomic element category, HMM chromatin states. A, UCSC genome browser representation of the region encompassing the preselected INHBB gene. B, Corre plots showing relevant features (see legends) for each one of the gene associated probes (anchor CpGs). C, Distribution of DNA methylation levels. D, Quantities of correlating sites in normal and tumor tissues. E, Genomic distribution of co-methylations, colored by chromosome.

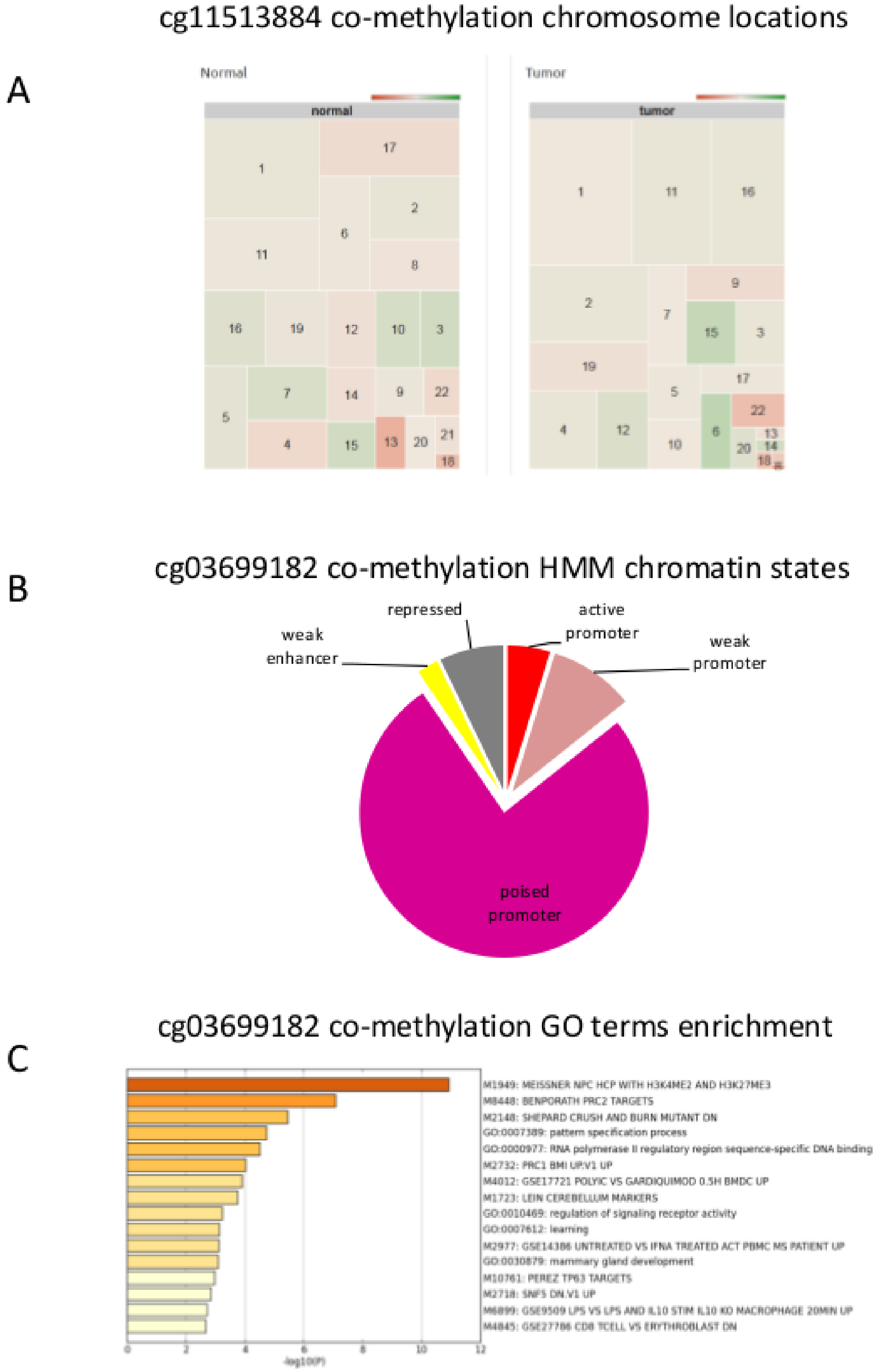

Figure S30: Corre usage example: INHBB. A, Treemap of chromosome distribution of co-methylating probes with cg11513884 located 20kb upstream of the INHBB gene (see Figure 7A). B, Distribution of HMM chromatin states in sites co-methylating with probe cg03699182 located in the INHBB gene CpG island promoter. C, Biological and functional enrichment of genes associated with the cg0369182 co-methylation probes.
